# BCR, not TCR, repertoire diversity is associated with favorable COVID-19 prognosis

**DOI:** 10.1101/2024.06.11.598368

**Authors:** Faith Jessica Paran, Rieko Oyama, Abdullah Khasawneh, Tomohiko Ai, Hendra Saputra Ismanto, Aalaa Alrahman Sherif, Dianita Susilo Saputri, Chikako Ono, Mizue Saita, Satomi Takei, Yuki Horiuchi, Ken Yagi, Matsuura DVM Yoshiharu, Yasushi Okazaki, Kazuhisa Takahashi, Daron M Standley, Yoko Tabe, Toshio Naito

## Abstract

The SARS-CoV-2 pandemic has had a widespread and severe impact on society, yet there have also been instances of remarkable recovery, even in critically ill patients. In this study, we used single-cell RNA sequencing to analyze the immune responses in recovered and deceased COVID-19 patients during moderate and critical stages. The study included three unvaccinated patients from each outcome category. Although expanded T cell receptor (TCR) clones were predominantly SARS-CoV-2-specific, they represented only a small fraction of the total repertoire in all patients. In contrast, while deceased patients exhibited monoclonal B cell receptor (BCR) expansions without COVID-19 specificity, survivors demonstrated diverse and specific BCR clones. These findings suggest that neither TCR diversity nor BCR monoclonal expansions are sufficient for viral clearance and subsequent recovery. Differential gene expression analysis revealed that protein biosynthetic processes were enriched in survivors, but that potentially damaging mitochondrial ATP metabolism was activated in the deceased. This study underscores that BCR repertoire diversity, but not TCR diversity, correlates with favorable outcomes in COVID-19.

## Introduction

Coronavirus disease 2019 (COVID-19), caused by severe acute respiratory syndrome coronavirus 2 (SARS-CoV-2), has infected more than 660 million people since late 2019. Symptoms of SARS-CoV-2 infection vary widely: some patients are asymptomatic, while others suffer fatal symptoms. A wealth of studies has shown that differential immune responses to infection can contribute to these wide-ranging outcomes^1, 2, 3, 4^.

Gene expression studies of severe COVID-19 have revealed that in peripheral blood mononuclear cells (PBMCs), interferon-stimulated genes (ISGs) were highly upregulated in most cell types, including T and B cells^5, 6^, but diminished as disease severity progressed^7^. In addition to the inflammatory responses, Luo and colleagues reported that pro-inflammatory T cell transcriptional profiles shifted toward metabolic adaptation in patients recovering from severe COVID-19^8^. Consistently, disruption of mitochondrial energy metabolism has been observed in severe COVID-19 cases^9, 10^. These findings indicate that metabolic adaptation and activation are important for survival of immune cells during COVID-19 infection, although the details of this process remain to be elucidated.

Adaptive immunity is mediated by repertoires of T cell receptors (TCRs) and B cell receptors (BCRs) expressed on the surfaces of T and B cells, respectively^11^. Due to somatic recombination of V(D)J genes, the potential diversity of adaptive immune receptor repertoires far exceeds the number of B or T cells in any one individual. In B cells, this diversity is further enhanced by the process of somatic hypermutation (SHM), which is crucial for increasing the affinity of BCRs to antigens, and introduces additional variability into the BCR repertoire^12^. As a result, only a small fraction of the BCRs and TCRs that can be observed in any one individual are likely to be observed in another individual, even when they both have been vaccinated at the same time^13^. Although adaptive immune repertoires are strongly shaped by the antigens they encounter, the diversity in any given donor makes it challenging to differentiate patients in terms of disease severity based only on observed repertoire data.

In the case of COVID-19, studies of T and B cell responses have been abundant, with 292 T cell-based studies and 132 B cell-based studies, amounting to millions of TCRs and BCRs in the public domain at the time of this writing (PubMed, accessed on 2023-12-04). However, interpretation of such data is at times conflicting. For instance, reduced TCR diversity in COVID-19 patients was reported to be associated with poor outcomes^14, 15^, while monoclonal expansion was reported to be essential for viral clearance by another study^16^.

Both expansion^17, 18^ and contraction^19, 20^ of CD4 and CD8 T cell clones in critically ill patients compared to mild COVID-19 cases have been reported. Analysis of B cell responses have been somewhat more consistent. For example, COVID-19 patients were reported to show greater monoclonal expansion of BCR compared to healthy controls^21^. BCRs with monoclonal expansion in COVID-19 patients have been reported to include the IGHV3 gene family (e.g., IGHV3-23 and IGHV3-30)^22, 23^ as well as IGHV1 and IGHV4 gene families^24^. However, BCR antigen specificity is mediated by specific sequences in the complementarity determining regions (CDRs), which are not solely defined by any single V gene, but rather influenced by SHM, which introduces additional variability into the BCR repertoire. Furthermore, the overall trends in BCR repertoires do not necessarily describe or predict the B cell responses in any one patient.

In addition to the considerations above, repertoire analysis in the post-pandemic era is complicated by the fact that most donors have encountered SARS-CoV-2 antigens, either through infection or vaccination. To isolate features of individual immune responses that are associated with disease severity and survival, it is advantageous to examine donors undergoing response to the virus for the first time.

To this end, we investigated adaptive immune responses in three groups of unvaccinated Japanese donors, including COVID-19 patients who ultimately recovered, patients who experienced fatal outcomes, and, as a control, healthy uninfected donors. Both gene expression, and TCR and BCR repertoires were profiled at the single cell level as a function of time. From the single cell gene expression data, we observed a shift in gene expression from inflammatory responses to biosynthetic processes or energy metabolism pathways, consistent with previous studies.

The repertoires of each donor were examined individually, at each time point, and between donors. In addition, we examined the individual repertoires in the context of the vast accumulated TCR and BCR data from other COVID-19 repertoire studies. From the individual TCR and BCR repertoires, we noted clear differences in diversity and monoclonal expansions in each individual, with fatal patients exhibiting relatively large monoclonal B cell expansions and surviving patients showing greater BCR diversity. Interestingly, when examined in the context of past COVID-19 studies we observed that the patients who ultimately recovered exhibited overlap in their repertoires with previously published COVID-19 patient repertoires to a significantly greater extent than did the fatal cases. Expression of monoclonally expanded BCRs in the fatal patients indicated that the expanded B cells were not specific to known SARS-CoV-2 antigens. Consistently, the proportion of BCR matches to COVID-19 specific BCRs were significantly higher in recovered patients than fatal patients.

## Results

In this study, we analyzed samples from six COVID-19 patients whose onset of symptoms ranged from November 13, 2020, to June 7, 2021. Three of the deceased patients (D 1-3) experienced fatal symptoms while the other three recovered (R 1-3). None of these patients had been vaccinated, allowing differential immune responses in the two groups to be analyzed without complications from prior exposure to SARS-CoV-2 antigens. During this period in Tokyo, where the samples were collected, the number of cases registered in GISAID EpiCoV database^25^ increased from 200 in October 2020 to 2,500 by July 2021. The predominant lineages shifted from B.1.1 to B.1.1.89 (accessed on 2023-11-16). All dead patients were infected with the B.1.1.7 subcluster of the Alpha strain during the fifth wave. Recovering patient, R1, was infected with the R.1. lineage (fourth wave) which was identified in the United States and Japan; R2 and R3 were both infected with the B.1.1.214 during the fifth wave. Detailed information about the patients is provided in Table 1.

**Table 1.**
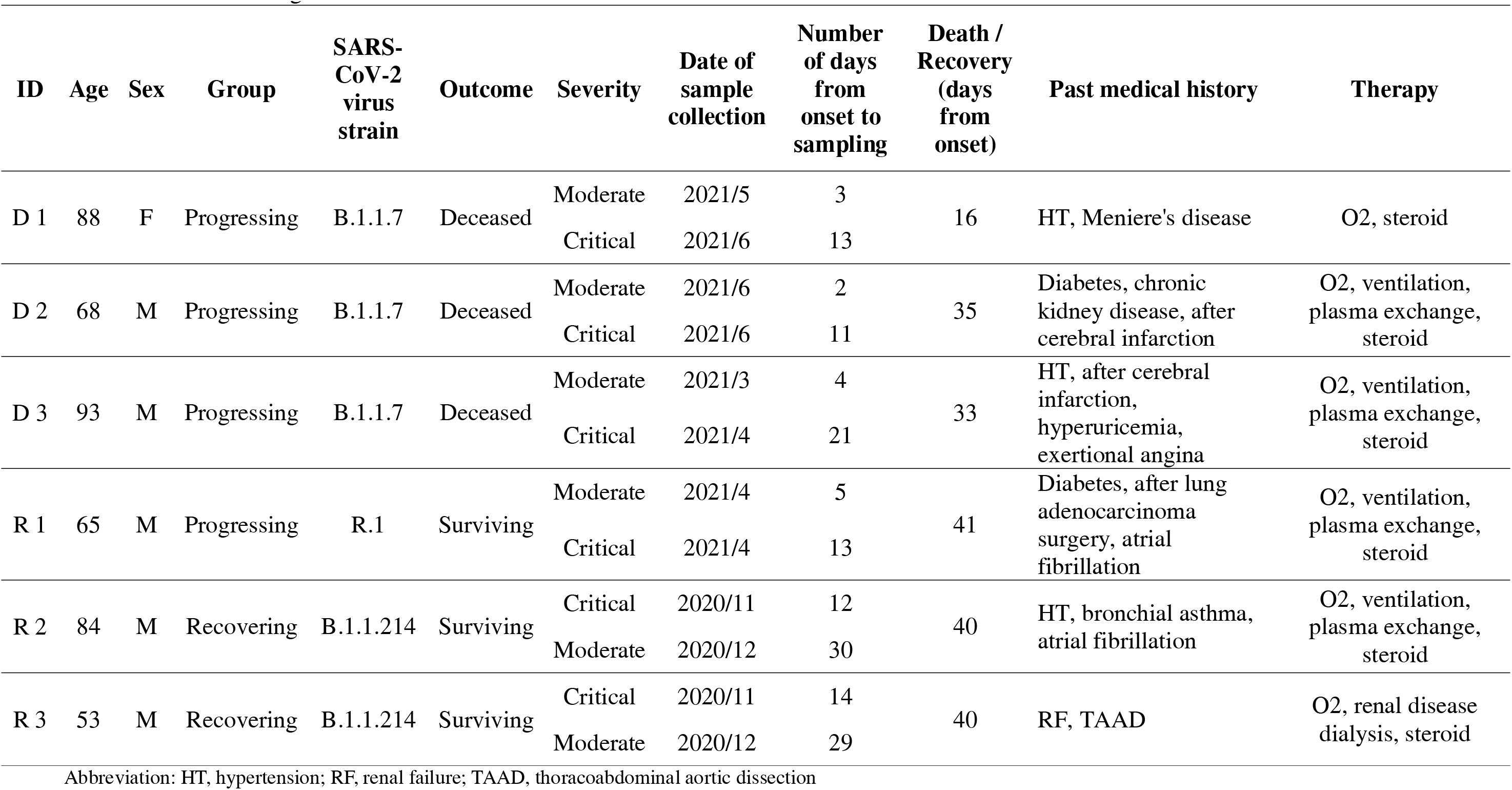
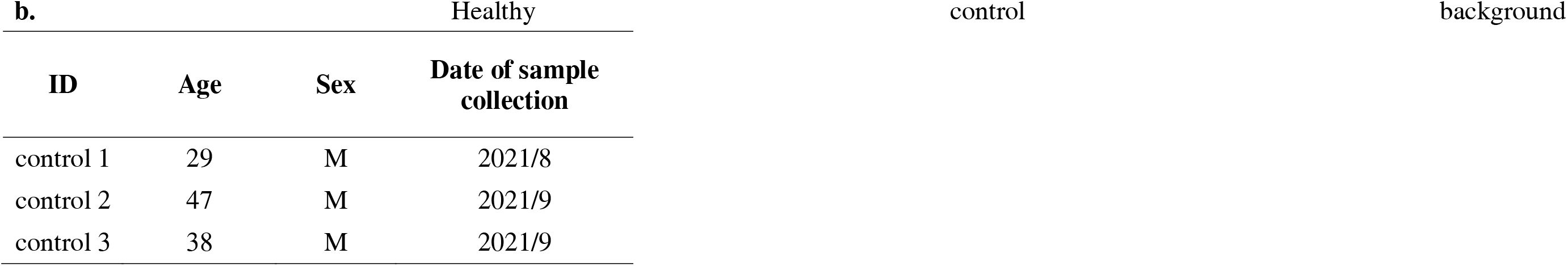
**a.** Patient background.

| ID | Age | Sex | Group | SARS-CoV-2 virus strain | Outcome | Severity | Date of sample collection | Number of days from onset to sampling | Death / Recovery (days from onset) | Past medical history | Therapy |
| --- | --- | --- | --- | --- | --- | --- | --- | --- | --- | --- | --- |
| D 1 | 88 | F | Progressing | B.1.1.7 | Deceased | Moderate | 2021/5 | 3 | 16 | HT, Meniere's disease | O2, steroid |
|  |  |  |  |  |  | Critical | 2021/6 | 13 |  |  |  |
| D 2 | 68 | M | Progressing | B.1.1.7 | Deceased | Moderate | 2021/6 | 2 | 35 | Diabetes, chronic kidney disease, after cerebral infarction | O2, ventilation, plasma exchange, steroid |
|  |  |  |  |  |  | Critical | 2021/6 | 11 |  |  |  |
| D 3 | 93 | M | Progressing | B.1.1.7 | Deceased | Moderate | 2021/3 | 4 | 33 | HT, after cerebral infarction, hyperuricemia, exertional angina | O2, ventilation, plasma exchange, steroid |
|  |  |  |  |  |  | Critical | 2021/4 | 21 |  |  |  |
| R 1 | 65 | M | Progressing | R.1 | Surviving | Moderate | 2021/4 | 5 | 41 | Diabetes, after lung adenocarcinoma surgery, atrial fibrillation | O2, ventilation, plasma exchange, steroid |
|  |  |  |  |  |  | Critical | 2021/4 | 13 |  |  |  |
| R 2 | 84 | M | Recovering | B.1.1.214 | Surviving | Critical | 2020/11 | 12 | 40 | HT, bronchial asthma, atrial fibrillation | O2, ventilation, plasma exchange, steroid |
|  |  |  |  |  |  | Moderate | 2020/12 | 30 |  |  |  |
| R 3 | 53 | M | Recovering | B.1.1.214 | Surviving | Critical | 2020/11 | 14 | 40 | RF, TAAD | O2, renal disease dialysis, steroid |
|  |  |  |  |  |  | Moderate | 2020/12 | 29 |  |  |  |

Abbreviation: HT, hypertension; RF, renal failure; TAAD, thoracoabdominal aortic dissection

| b. Healthy |  |  |  | control | background |
| --- | --- | --- | --- | --- | --- |
| ID | Age | Sex | Date of sample collection |  |  |
| control 1 | 29 | M | 2021/8 |  |  |
| control 2 | 47 | M | 2021/9 |  |  |
| control 3 | 38 | M | 2021/9 |  |  |

A total of 129,469 cells were obtained for downstream analyses, and thirty cell clusters were identified and then annotated by the Azimuth web application (Figure 1). Of these, 30,071 (23.2%) were from healthy subjects, 47,657 (36.8%) were from samples in the moderate stage, and 51,723 (40%) were from samples in the critical stage. Fine resolution clustering of mRNA profiles revealed 13 clusters of T lymphocytes, annotated as follows: CD4 and regulatory T (Treg) subtypes, CD8 subtypes, mucosal associated invariant T (MAIT) cells, double-negative T cells (dnT), and gamma-delta T cells (gdT). The detected B cell subtypes were B naïve, B intermediate, B memory, and plasmablasts.

**Figure 1.**
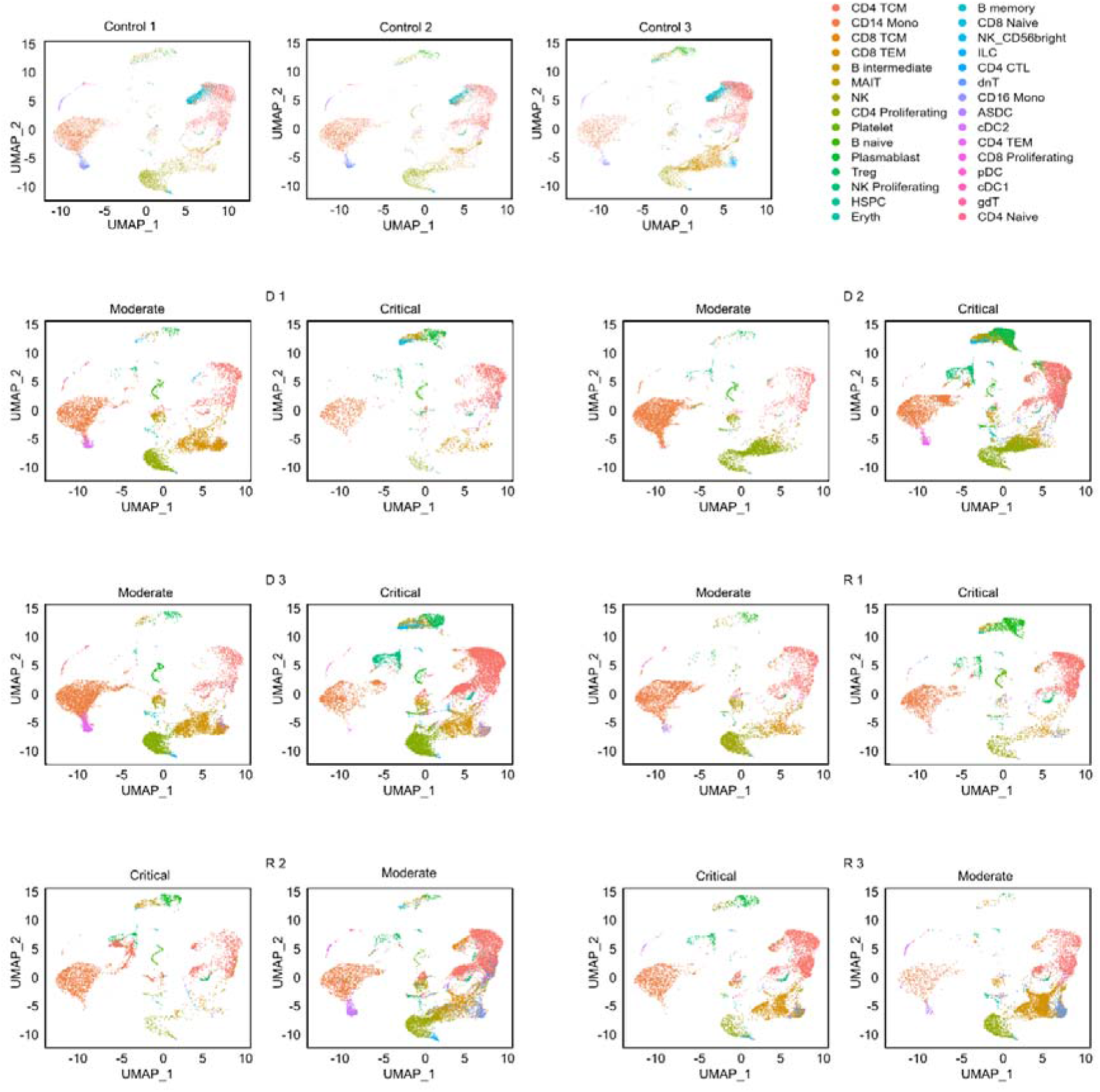
UMAP projections of cell populations in the healthy and COVID-19 samples. Azimuth annotation of clustered peripheral mononuclear blood cells (PBMCs) shown through UMAP projection shows the general trend of cell proportions with relation to disease severity. Legend: CD4 CTL, CD4-positive cytotoxic T lymphocytes; CD4/CD8 TEM, CD4-/CD8-positive effector memory T cell; CD4/CD8 TCM, CD4-/CD8-positive central memory T cell; Treg, regulatory T cell; MAIT, Mucosal associated invariant T cell; NK, natural killer cell; Eryth, erythroid cell; HSPC, hematopoietic precursor cell; ASDC, AXL+ dendritic cell; ILC, innate lymphoid cell; cDC1, CD141-positive myeloid dendritic cell; cDC2, CD1c-positive myeloid dendritic cell; pDC, plasmacytoid dendritic cell.

### Populations of T cell subtypes

A total of 61,327 T cells from all patients and healthy subjects were recovered after merging and filtering the single-cell RNA sequencing (scRNA-seq) dataset with the TCR assembly (Table S1). Naïve T cells were significantly reduced in all six patients compared to the healthy subjects (Figure 2; Table S1), indicating activation of the adaptive immune response upon infection. The most abundant T cell subtype was CD4 memory (CD4 TCM) with 32,262 cells, followed by CD8 effector (CD8 TEM) with 18,111 cells. All other T cell subtypes each had fewer than 3,000 cells. Therefore, downstream analyses were focused on the CD4 TCM and CD8 TEM subsets.

**Figure 2.**
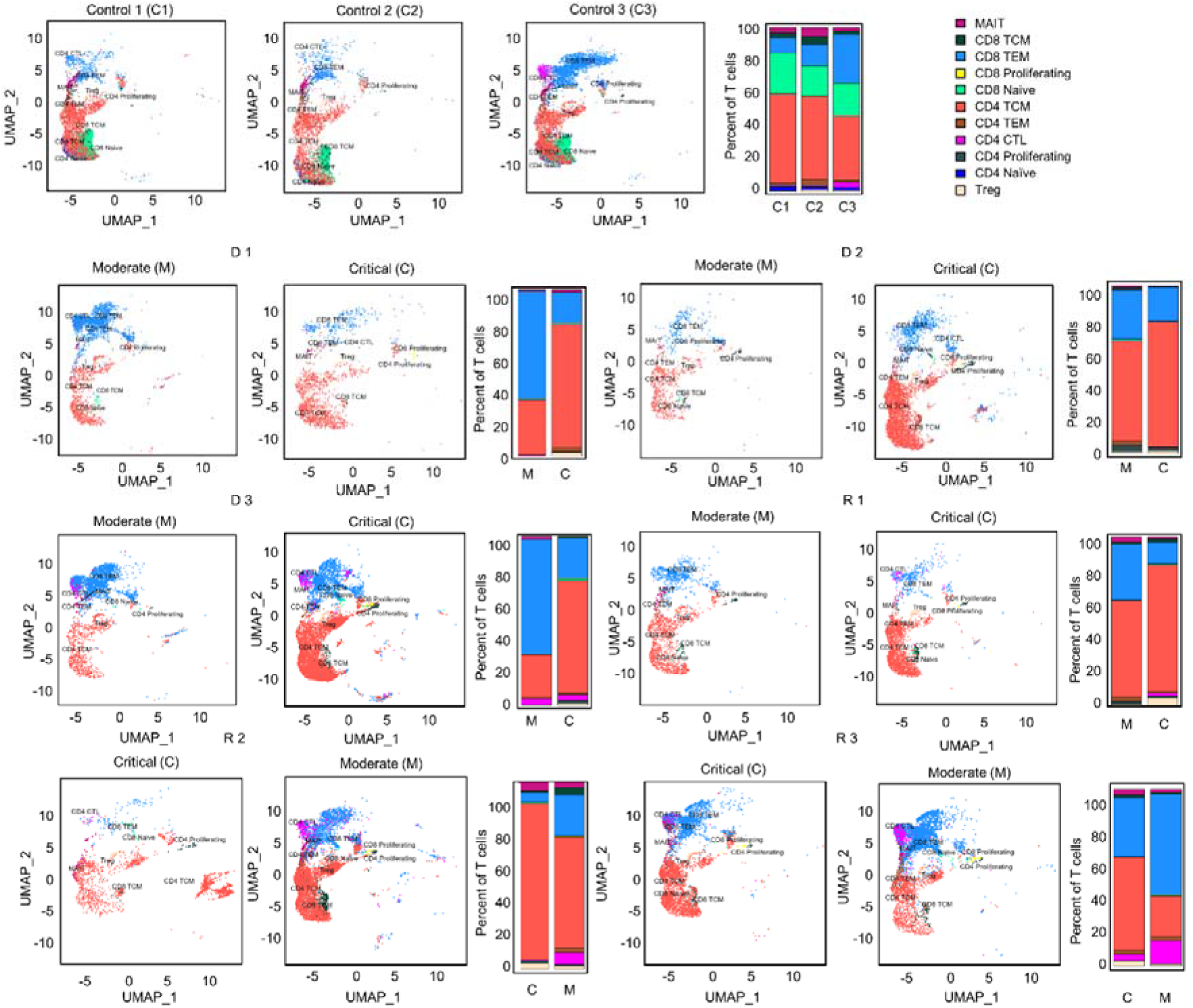
UMAP projections and bar graphs for T cell clusters and proportions. Eleven clusters were generated for T cells. Bar plot shows cell compositions at moderate and critical stages, with CD4 TCM and CD8 TEM occupying most of the T cells for healthy subjects and COVID-19 patients. Healthy subjects had a high proportion of CD8 naïve cells. Legend: CD4 CTL, CD4-positive cytotoxic T lymphocytes; CD4/CD8 TEM, CD4-/CD8-positive effector memory T cell; CD4/CD8 TCM, CD4-/CD8-positive central memory T cell; Treg, regulatory T cell; MAIT, Mucosal associated invariant T cell.

The CD4 TCM cell ratio significantly increased as the disease progressed from moderate to critical in D 1-3 and R 1 patients (Mann-Whitney U test, *p* < 0.05), but decreased as R 2 and 3 recovered from critical to moderate—although not significantly. Proportions of CD8 TEM cells were reduced from moderate to critical in deteriorating patients D 1-3 and R 1 (Mann-Whitney U test, *p* < 0.05), except D 2 whose CD8 TEM proportions were stable. In contrast, CD8 TEM cells of R 2 and 3 increased as they recovered from the critical state to moderate.

### Inflammatory genes in T cells were highly expressed in early moderate stages

To determine the changes in each patient’s transcriptome throughout the duration of the disease, we compared differences in gene expression between moderate and critical stages (threshold parameters: average log2FC > 0.58; Bonferroni adjusted *p < 0.05*). All patients progressed to critical 11–14 days from onset. The moderate stages were of two different timepoints. D 1-3 and R 1 were sampled during the early stages of the disease (2-5 days from onset) before progressing to the critical state. R 2 and 3, however, were sampled during the recovery phase (41 days and 29 days after onset, respectively).

Expanded T cell receptors are known to respond to specific non-self antigens. Consequently, we performed differential gene expression analysis only on T cells expressing abundant V-gene regions in CD4 TCM cells (TRBV20-1, TRBV5-1, and TRBV19) to investigate whether these expanded receptors are involved in distinct different biological functions.

The expression patterns shown in Figure 3a differed significantly between moderate and critical conditions for progressing D 1-3 and R 1, where inflammatory genes, such as *IFI44L, IFITM1, IFI6, ISG15, XAF1, STAT1, LY6E, B2M,* and *MX1* were highly expressed during the early moderate stage, while ribosomal and elongation factor genes (*RPS, RPL,* and *EEF1*) were dominant in the critical state. In contrast, in the critical states of R 2 and 3, genes involved in cellular respiration and energy transport, along with genes expressing ribosomal proteins in the moderate compared to the critical stage were upregulated. As R 2 and 3 recovered to moderate, fewer genes were differentially expressed, but mitochondrial genes (*MT-CO1, MT-CYB, MT-ND4L*) were prominent. The complete list of differentially expressed genes for each TRBV subset is summarized in Table S2.

**Figure 3.**
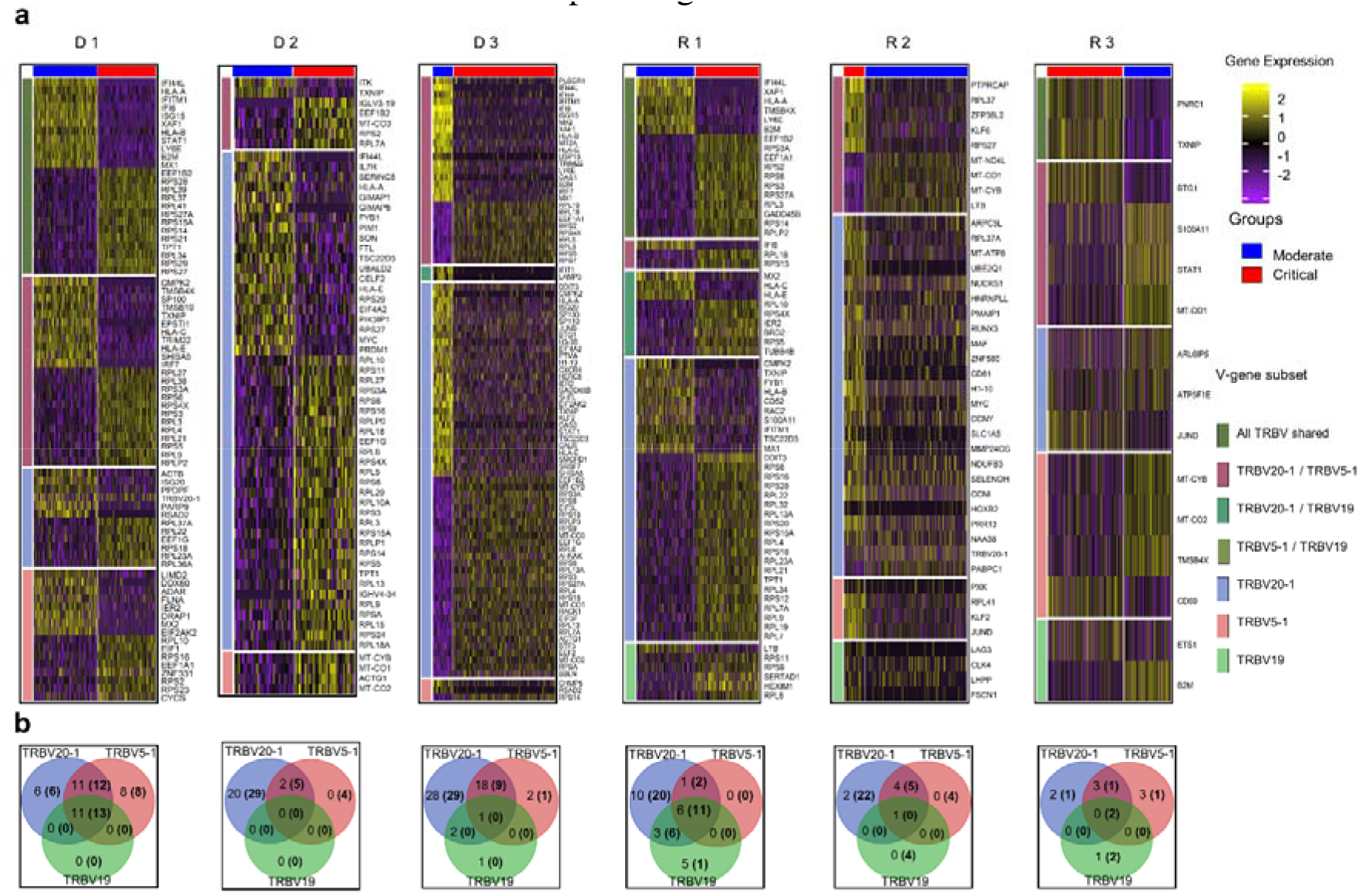
Differential gene expression analysis in CD4 TCM cells. **(a)** Normalized expression values of differential genes during the moderate and critical states for each patient are visualized in the heatmap. Inflammatory genes were upregulated in the moderate state of progressing patients (D 1-3 and R 1). Horizontal divisions represent the genes expressed and overlapped in the TRBV subsets. **(b)** Gene overlap among TRBV subsets upregulated and downregulated (in parentheses) during the moderate state, in comparison to the patient’s critical state. Genes in the non-intersection areas were defined as the specific gene sets for each TRBV subsets.

To further understand the differences in biological processes and cellular functions of the TRBV-gene regions, we performed gene overlap analysis of the three TRBV subsets for each patient (Figure 3b) using the genes recovered in Figure 3a. The TRBV20-1 gene was expressed more highly than TRBV5-1 or TRBV19. The differential genes expressed in TRBV5-1 and TRBV19 subsets were mostly shared with other TRBV subsets. TRBV20-1 had the greatest number of exclusive genes expressed in almost all patients. The exclusive genes from TRBV20-1 of progressing patients D 1-3 and R 1 were mostly inflammatory genes, and the downregulated genes were ribosomal and elongation factor genes. Shared and exclusive genes in all three TRBV subsets were much fewer in recovering patients R 2 and 3 during the critical and moderate states.

To determine the difference in activated biological pathways between the early stage of the disease (progressing group) and convalescing stage (recovering group), we performed gene set enrichment analysis (GSEA) of functional hallmark gene sets using the recovered genes from the previous analysis, and plotted the pathways activated in the moderate stage (Figure 4a, Table S3). In general, immune response pathways were enriched both in the early stage and convalescing stage of the disease. At the same time, pathways involved in proliferation and signaling were downregulated. This pattern was observed in all TRBV subsets, and no preference for specific gene families was found.

**Figure 4.**
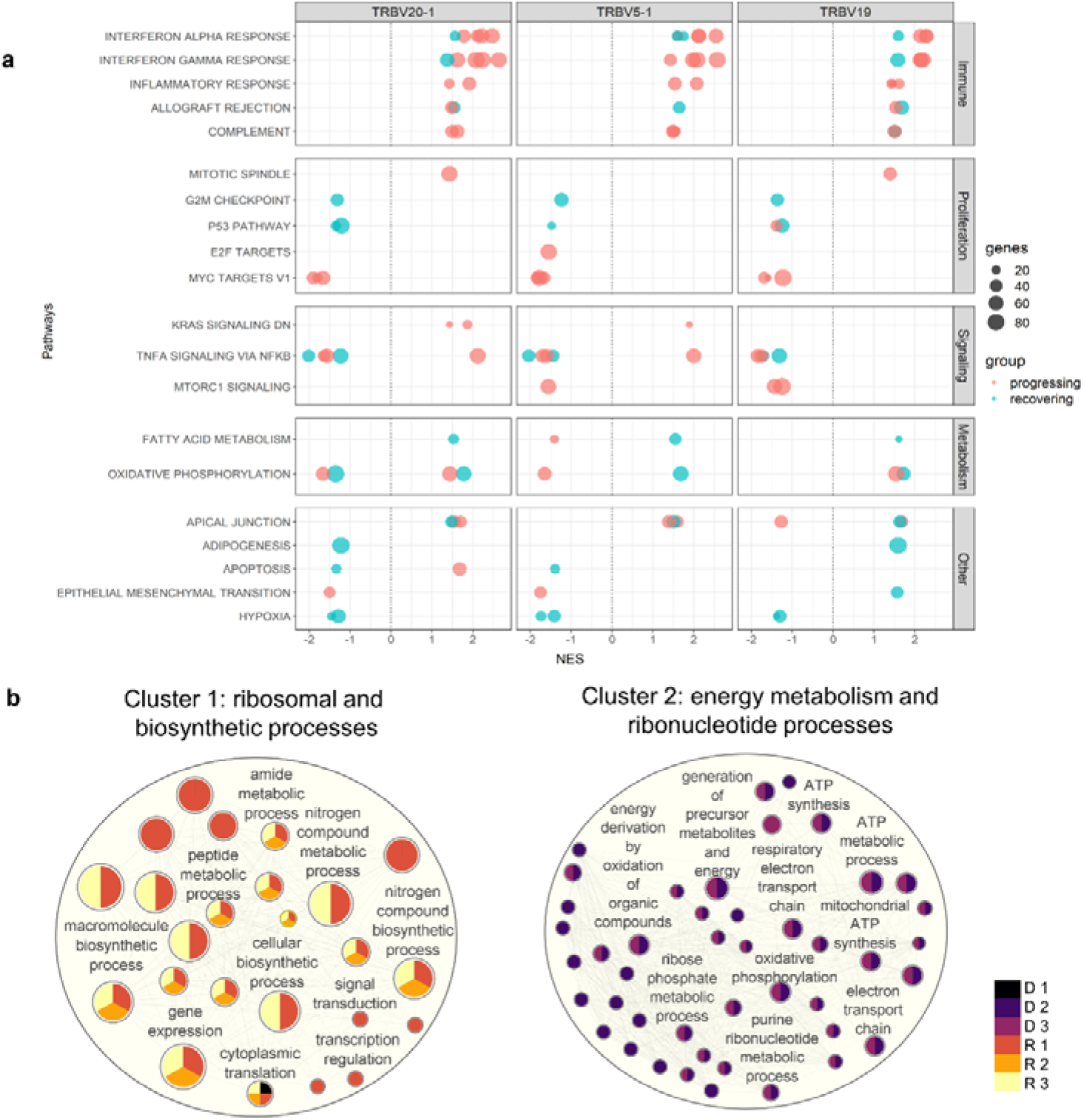
Pathway enrichment analysis in CD4 TCM cells. **(a)** Gene set enrichment analysis was performed between the moderate vs critical state of each patient. Plot summarized all Hallmark pathways activated and suppressed during the moderate states of progressing and recovering patients, divided by pathway category. Panels represent TRBV subsets. Red dots indicate progressing patients (D 1-3), and blue dots indicate recovering patients (R 1-3). **(b)** Gene ontology enrichment analysis was performed in go:Profiler using differentially expressed genes identified during the critical stage of deceased vs surviving patients. Nodes represent pathways; node colors represent patients; node sizes are proportional to number of genes associated with each pathway; edges represent connections between pathways. The clusters, calculated using MCL algorithm, represent similarities among the pathways.

Consistent with the differential gene expression, Interferon-α and Interferon-γ responses were the most enriched pathways during the moderate state of progressing patients in the TRBV20-1, TRBV5-1, and TRBV19 subsets. In contrast, *MYC* targets regulating several receptor-signaling functions of virus-infected host cells were highly downregulated in the moderate state of progressing patients.

While the interferon pathways were also enriched during the moderate state of recovery (R 2 and 3), enrichment scores were lower compared to the progressing patients (D 1-3 and R 1); only *B2M* (log2FC < 0.80) and *STAT1* (log2FC < 1.30) inflammatory genes were differentially expressed. Metabolic processes such as fatty acid metabolism and oxidative phosphorylation were activated in all three TRBV subsets for R 2 and 3, except for suppression for R 2 in TRBV20-1. Proliferation and signaling pathways, as well as hypoxia, were downregulated in R 2 and 4. Progressing patients during the moderate state had a higher enrichment score for interferon responses compared to recovering patients.

### Surviving patients exhibited active protein synthesis during critical state

Since TRBV5-1 and TRBV19 showed redundant CD4 TCM functions based on gene expression analysis, only cells with TRBV20-1 were used in this analysis. To determine the differences in biological processes activated during the critical state of deceased and surviving patients, we generated an enrichment network of the biological processes in Cytoscape^26^. Using the Markov Cluster (MCL) algorithm, two major groups of pathways, Cluster 1 and 2, were detected (Figure 4b). Cluster 1 was characterized by 25 pathways involved in protein metabolic and biosynthetic processes and was enriched in surviving patients (R 1-3); Cluster 2 was characterized by 44 pathways involved in mitochondrial energy metabolism processes and was enriched in deceased patients (D 2-3). However, deceased patient D 1 was grouped in the first cluster with the cytoplasmic translation pathway significantly activated during the critical state associated with ribosomal protein-coding genes *RPL17, RPS29, RPS4X,* and *RPL38*. The complete lists of DEGs and pathways recovered for the enrichment network are shown in Table S4 and Table S5, respectively.

We next utilized the unique molecular ID (UMI) counts to assess the transcriptional activity of cell clusters. High UMI counts represent dynamic transcriptional status that could cause replication stress^9^. During the critical phase, high UMI values were observed in the CD4 proliferating cell cluster of the deceased patients (normalized UMI: deceased D 1 = 31.2; D 2 = 199.8; D 3 = 226.1; surviving R 1 = 27.9; R 2 = 51.1; R 3 = 41.0). The UMI values were positively associated with cell counts (CD4 Proliferating cell count and proportion in parentheses during the critical state: deceased D 1 = 2 (0.1); D 2 = 82 (1.9); D 3 = 134 (1.6); surviving R 1 = 26 (1); R 2 = 30 (1.4); R 3 = 13 (0.3)). Notably, only the pro-inflammatory gene *LTB,* implicated in “cytokine storm” ^27, 28^, and *MT-ATP6* were significantly upregulated in the CD4 proliferating cells of deceased patients compared to survivors (avg_log2FC: *LTB* 0.9, *MT-ATP6* 1.12) (list of DEGs in Table S6).

These findings suggest that during the critical phase, the overexpression of mitochondrial genes involved in energy production and the increase in inflammatory signatures in CD4 proliferating cells may indicate an unfavorable outcome.

### Diverse COVID-19 specific TCR repertoires were observed in all patient groups

To predict the general reactivity of T cell receptors to SARS-CoV-2, the CDR3 region of the TCRβ chain was cross-referenced to four public TCR databases (McPAS^29^, vdjdb^30^, TCRex^31^, and TCRMatch^32^).

Of the 44,486 filtered clonotypes identified for the TCR repertoire from TCR sequencing, 6,185 CDR3β sequences matched with 97-100% sequence similarity to one or more reference TCRs with a known viral antigen. Of these, over half (3,688) showed high specificity to a known SARS-CoV-2 antigen (Table S2), representing 8.3% of the total TCR repertoire. The proportion of unique clonotypes matching to viral antigens was expressed as a percentage of the total TCR repertoire; the proportion of clonotypes matching to SAR-CoV-2 antigens did not differ among donors, including healthy subjects. Whether this observation represents true SARS-CoV-2-specific TCR clonotypes present in the unvaccinated healthy donors or simply reflects a lack of well-annotated TCR sequences remains an open question.

We next focused on the most abundant clonotypes, which we assumed would be relevant to host defense against the virus. We investigated the occupancy of the top 50 clonotypes to account for the diversity in the entire TCR repertoire. We ranked the clonotypes by abundance during the critical state and plotted the cumulative frequency for each sample (Figure S1). In all cases, multiple clonotypes predicted to target COVID-19 peptide antigen (red dots) were present in the top clonotypes. The mean occupancy of the top 50 clonotypes was significantly higher in COVID-19 cases (mean = 0.43%) than in healthy subjects (mean 0.20%) (unpaired T test, p < 0.05). However, when comparing the percent occupancy of all clonotypes in the TCR repertoire per critical sample, no significant difference was found among subjects, revealing comparable TCR repertoire diversity in all patients during the critical state.

The V-J gene usage preferences of the TCR repertoire can also be used to describe the TCR response to infection in relation to its overall diversity. We assessed the V-J gene segments in the TCR β-chain of the CD4 TCM and CD8 TEM cell subtypes. The most abundant V-J pair in CD4 TCM cells for both healthy subjects and patients included TRBV20-1, TRBV5-1, TRBV28, and TRBV19, in conjunction with TRBJ2-1, TRBJ2-3, TRBJ2-5, and TRBJ2-7 (Figure S2a; Table S8). The appearance of these abundant V-J gene regions in both healthy subjects and patients suggests their existence in the general population before the COVID-19 pandemic.

For CD8 TEM, the V-J usage was heterogeneous for each patient and had less diversity (Figure S2b; Table S9). Healthy subjects had higher diversity in V-J usage compared to the patients. The patients had monoclonal expansions of V segments and fewer combinations with J segments. For example, hyper expansions (cells > 100) were observed in the moderate stage of two deceased patients (D 1: TRBV4-1 and D 3: TRBV18), followed by drastic cell reduction in the critical stage. Hyper expansions were also observed in recovering patients (R 2-3), as well as shared V segments (e.g., TRBV28, TRBV7-9, and TRBV20-1), but mostly during the recovering phase. No systematic differences between patients and healthy subjects were observed in the CD8 TEM cells.

### Populations of B cell subtypes

Integration of the V(D)J sequences into the scRNA-seq data and filtering out of cells without heavy and/or light chain sequences yielded a total of 6,662 B cells from all patients and healthy subjects. These contained 2,260 Plasmablasts (33.9%), 2,326 B naïve (34.9%), 1,427 B intermediate (21.4%), and 649 B memory (9.7%) cells (Table S10). Comparisons of B cell proportions (Figure 5) showed high frequency of plasmablasts in COVID-19 patients compared to healthy subjects, but intermediate B cells were fewer in COVID-19 patients. A significant increase in plasmablasts was observed in the critical phase of deceased patients D 2-3 compared to all other patients. Given the wide variance in clonotype counts across different B cell subtypes, we took a comprehensive approach for subsequent analysis and assessed BCR diversity without segregating into subclasses.

**Figure 5.**
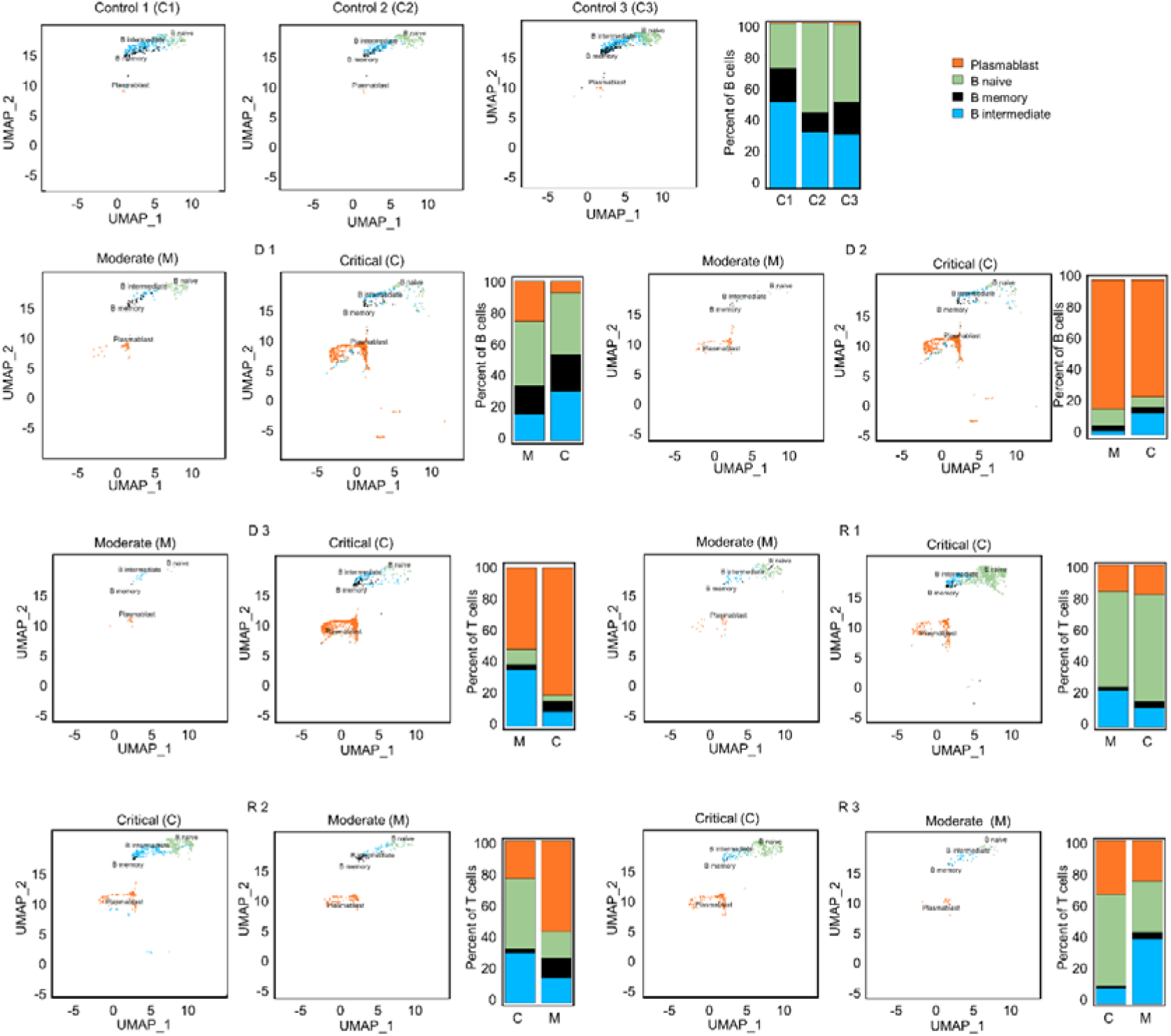
UMAP projections and bar graphs for B cell clusters and proportions. Four clusters were generated for B cells. Bar plot shows cell compositions at moderate and critical stages, with plasmablasts making up the majority B cells in deceased patients, D 2 and 3. In surviving patients, a larger proportion of naïve B cells is observed.

### Differentially expressed genes and GSEA in B cells

To determine the difference in activated biological pathways between the patients who deteriorated from moderate to critical disease (progressing group) and cases undergoing recovery from critical to moderate stages (recovering group), we performed GSEA of functional hallmark gene sets based on differentially expressed genes between the moderate and the critical samples for each patient.

In the progressing group (D 1-3 and R 1), upregulation of inflammatory genes such as *STAT1, IFITM1, LY6E,* and *ISG20* was predominant in the moderate stage before reaching the critical stage, and ribosomal protein (RP) genes were prominent in the critical stage (D 1-3 and R 1; Figure 6, Table S11). In the progressing group, enrichment scores for the Interferon Alpha/Gamma Response pathways were higher than in the recovering patients (Figure 7). In recovering patients (R 2 and 3), inflammatory gene expression decreased, and metabolic and biosynthetic gene expression increased in the moderate stage. Here, *MTORC1* signalling, fatty acid metabolism and oxidative phosphorylation were enriched in the moderate stage (Figure 6 and 7; Table S11 and S12). In all patients, activation of immune pathways and suppression of cell proliferation pathways were observed in the moderate state compared to the critical (Figure 7).

**Figure 6.**
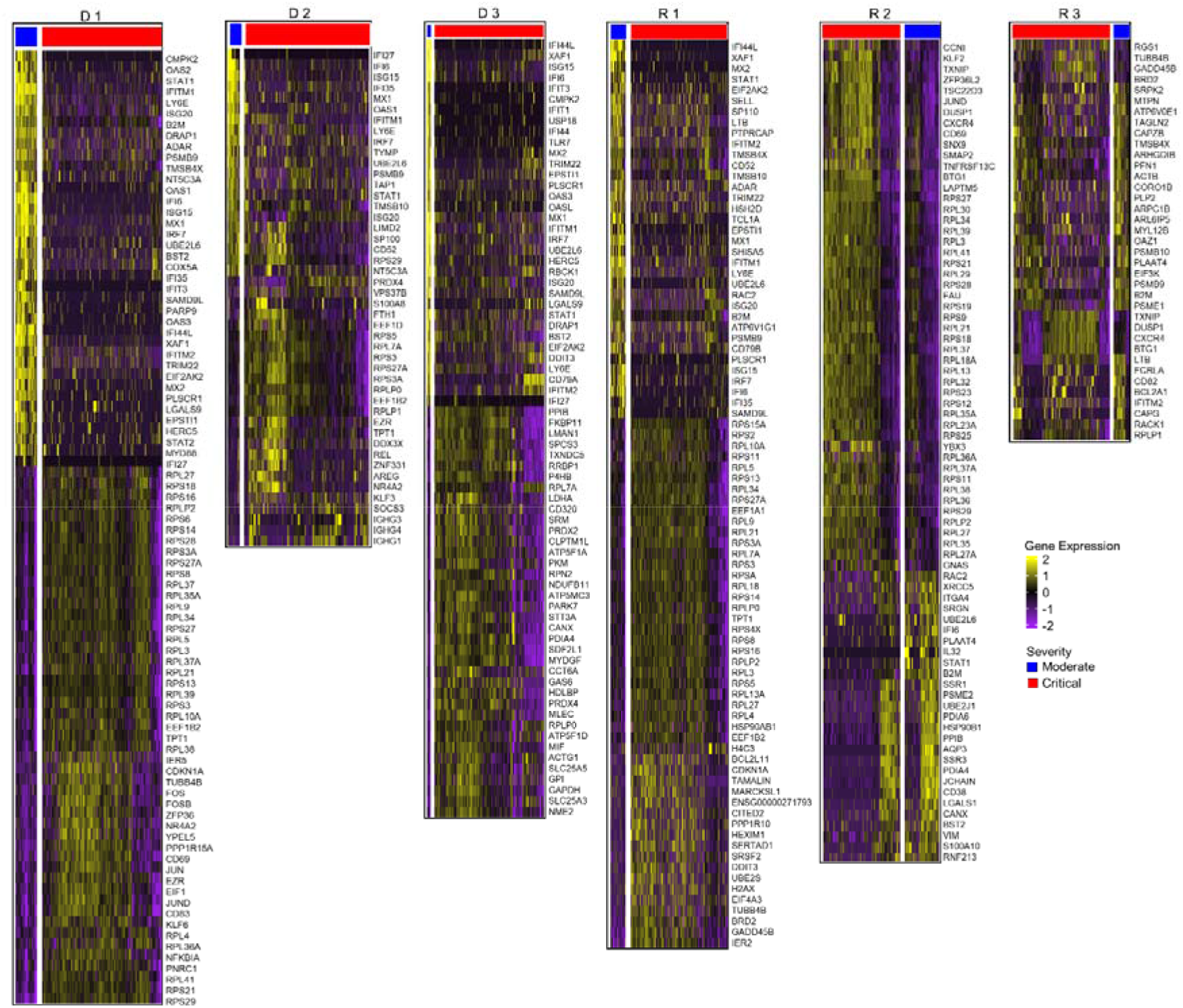
Differential gene expression analysis in B cells. Normalized expression values of differential genes during the moderate and critical states for each patient are visualized in the heatmaps. Patients who progressed in severity (D 1-3 and R 1) show upregulated inflammatory genes in their moderate, compared to the critical samples, where ribosomal protein genes were prominent. During the moderate state of recovering patients, R 2 and 3, metabolic and biosynthetic genes were upregulated. In critical samples, ribosomal proteins were upregulated.

**Figure 7.**
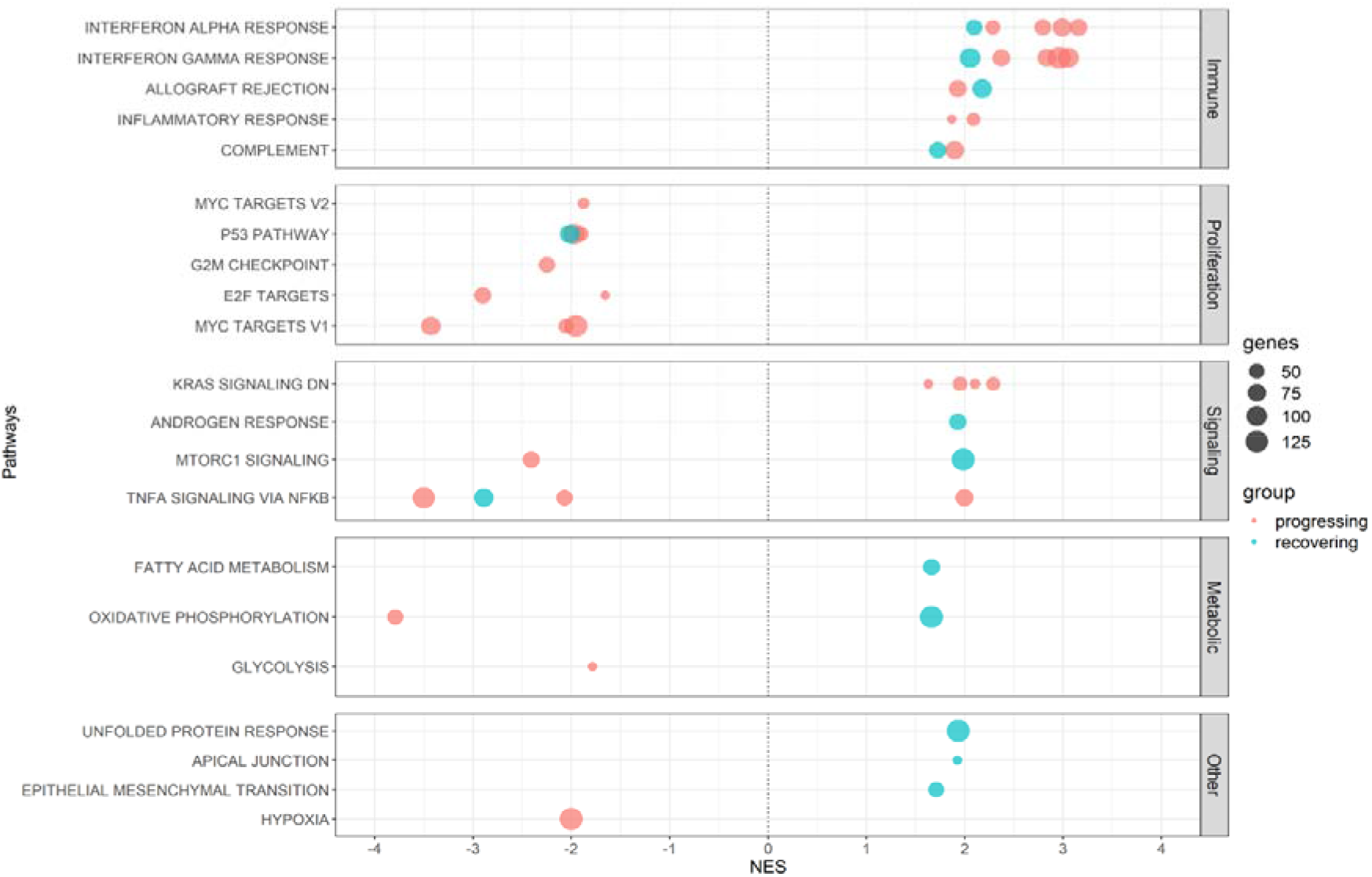
Shared enriched pathways in the moderate stages in B cells. Plot summarizes all Hallmark pathways activated and suppressed during the moderate states of progressing and recovering patients. Red dots indicate progressing patients (D 1-3), and blue dots indicate recovering patients (R 1-3). All patients had enriched immune pathways in this stage, with progressing patients having higher enrichment scores for the IFN-α and IFN-γ pathways. All patients had downregulated proliferation pathways.

We also performed GO analysis to compare differentially expressed genes of deceased and surviving patients during the critical stage. Oxidative phosphorylation, mitochondrial ATP synthesis, and various metabolic pathways were upregulated in the deceased patients. On the other hand, peptide and amide biosynthesis and metabolic pathways were upregulated in surviving patients (Table S13).

### BCR V-J gene usage

V(D)J recombination contributes to BCR diversity during the early stages of B cell maturation process, playing a key role in the formation of paratopes. Frequencies of V gene usage and proportions in B cells also change in response to infection (Table S14 and Figure S3). We investigated the variability in BCR V gene usage using the Shannon index. The range of V gene usage was significantly lower in COVID-19 patients than in healthy controls (*p* = 0.0029). The proportion of expanded clones (3 cells or more) was significantly higher in critical than in moderate phases (*p* = 0.008).

### Monoclonal expansion of BCR without SARS-CoV-2 affinity in deceased patients

Post V(D)J recombination, B cells undergo clonal selection and expansion, leading to the formation of greater BCR clonotypes. Because a higher frequency of clonotypes was observed during the critical stage than in the moderate stage across all patients (Table S10, S14), we focused on the heavy chain transcripts of the BCRs in the critical stage. The clonotypes were selected fully annotated BCR amino acid sequences for our study. The clonotype sequence counts for critical-stage COVID-19 samples in this assay were as follows: D 1 = 679, D 2 = 316, D 3 = 700, R 1 = 822, R 2 = 441, and R 3 = 398. Clonal expansion related to V-gene usage is represented by bubble plots (Figure 8a). In fatal patients, notable clonal expansions were observed in IGHV1-18 (cell count = 48, denoted as D1_1 in D 1), IGHV4-34 (cell count = 204, as D2_1 in D 2). Conversely, in recovered patients, less pronounced proliferations were noted with IGHV3-30 (cell count = 3, as R2_1 in R 2), and IGHV4-59 (cell count = 6, as R3_1 in R 3) showing fewer expansions.

**Figure 8.**
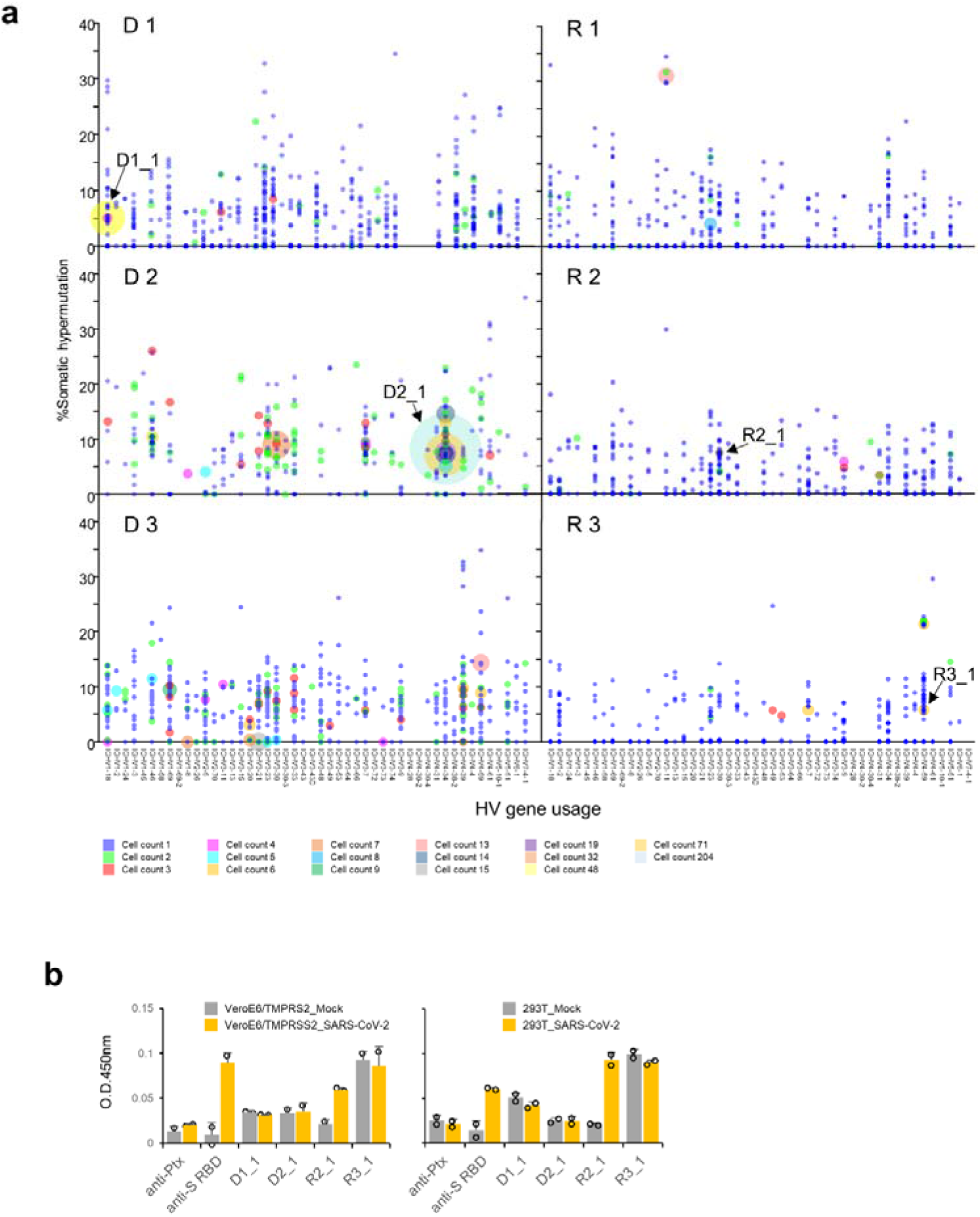
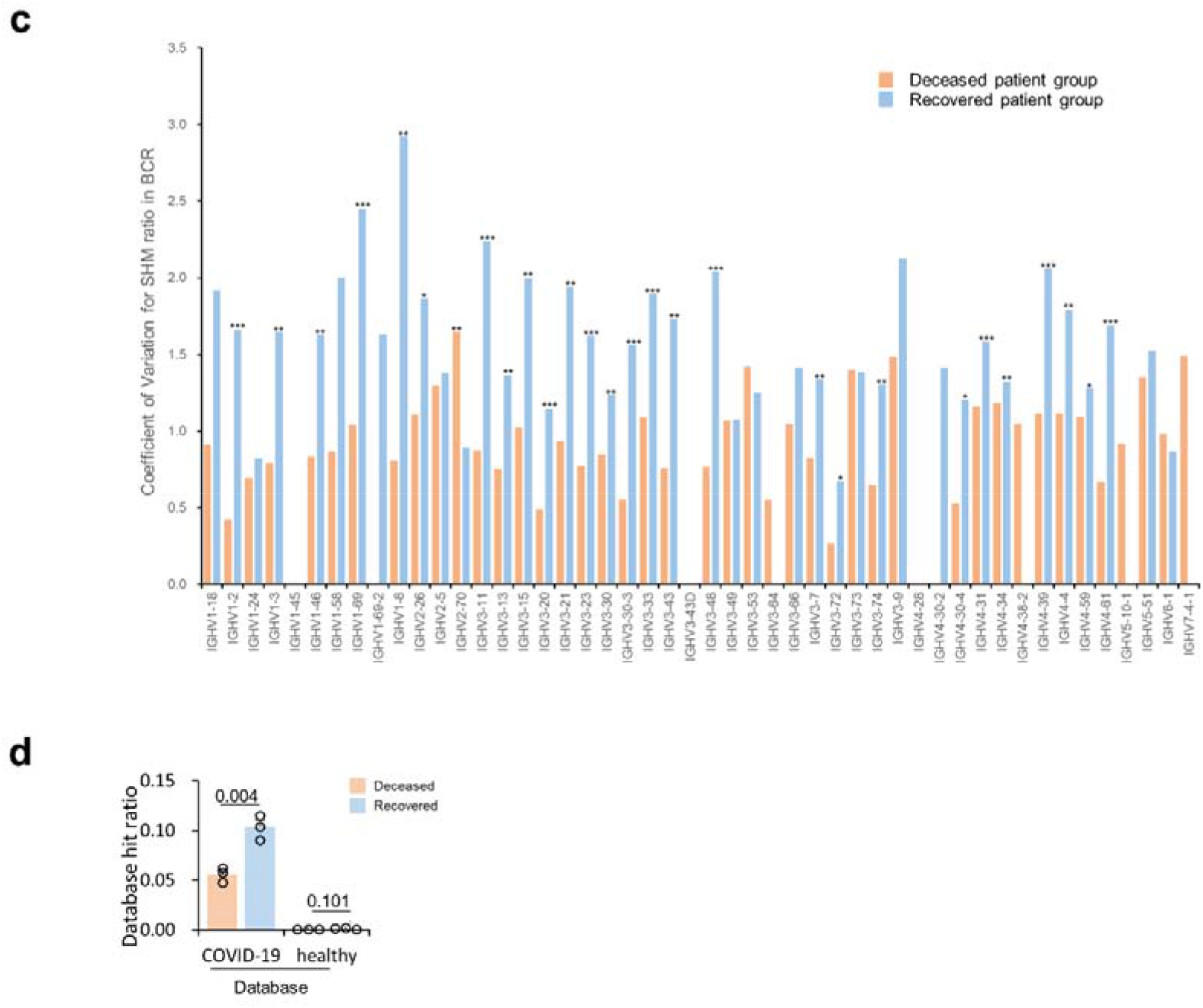
Detailed Verification of BCRs in COVID-19 Patients. **(a)** Somatic hypermutation and clonal expansion in critical samples. Individual clonotypes are depicted as bubbles with the size indicating the count of each clonotype and their position corresponding to HV gene usage and somatic hypermutation (SHM) rate. Larger bubbles denote higher clonotype counts. Clonal expansions are marked in the deceased patient groups (D 1-3) with counts exceeding 15. The recovered patient groups (R 1-3) show counts not exceeding 14. Arrows indicate the expanded clonotypes used for subsequent antibody binding assays. **(b)** Recombinant antibody binding to SARS-CoV-2 using expanded clones from COVID-19 patients. Variable region sequences from expanded clonotypes were expressed with the human IgG1 or kappa constant regions. The binding of these recombinant antibodies to immobilized SARS-CoV-2 infected call lysates was assessed using ELISA format. **(c)** Average coefficient of variation for somatic hypermutation (SHM) ratio in BCR from COVID-19 patients in the critical stage across different IGHV genes. Clone sequences were analyzed for SHM using IMGT/HighV-Quest. Statistical significance (determined by the Mann-Whitney U-test) between the groups for each HV-gene is indicated by the following markers: ***: p < 0.001, **: p < 0.05, *: p < 0.1. **(d)** Database hit ratios of CDR sequences between deceased and recovered patient groups. CDR1, CDR2, and CDR3 amino acid sequences from patient-derived clones were compared to a sequence database from prior COVID-19 studies and healthy individuals. Ratios are calculated as the number of database hits per total clonotypes in each patient’s heavy and light chains. The recovered patient group exhibited a significantly higher hit ratio compared to the deceased group (one-tailed Welch’s t-test, *p* = 0.0043).

Monoclonally expanded cells are reported to have the ability to respond to specific antigens^33^. We expressed recombinant antibodies from the clonally expanded BCR sequences of patients who recovered (R 2 and 3) and from patients who did not recover (D 1 and 2, the clonotypes are indicated in Figure 8a and S4). However, none of the expanded BCR clones in fatal patients recognized SARS-CoV-2 antigens in this assay. On the other hand, one antibody specific to SARS-CoV-2-infected cell lysates from recovered R 2 was observed (R2_1, as seen in both VeroE6/TMPRSS2 and 293T cells, in Figure 8b and S5). Based on these findings, we tentatively conclude that the BCR expansions in the fatal group of patients were not driven by affinity to known SARS-CoV-2 antigens.

### Higher coefficient of variation of SHM in recovered patients

Subsequent to V(D)J recombination, B cell maturation involves iterative cycles of SHM and selection dependent on antigen affinity. SHM results in amino acid changes in the CDRs, altering the structural conformation of BCRs and potentially impacting their affinity towards specific antigens. Consequently, even within similar V gene usage, diverse antigen affinities can arise due to these mutations. This process refines and diversifies the BCR repertoire, further enhancing the body’s ability to recognize and respond to a wide array of antigens. SHMs were assessed using the IMGT/HighV-QUEST program^34^ (version 3.6.1, reference directory released on 2023-1-30, analysis date 2023-10-3). The average number of nucleotide mutations within the variable region for each clonotype was higher in the deceased patient group than in the recovered group (one-tailed Welch’s t-test, *p* = 0.041, Figure S6a).

Additionally, a greater proportion of clonotypes with mutated nucleotides was also presented in the deceased patient group (*p* = 0.016, Figure S6b). Notably, in D 3, five clonotypes exhibited clonal expansions exceeding four counts, all without mutations (at IGHV1-18, IGHV1-8, IGHV3-21, IGHV3-23, and IGHV3-73, shown in Figure 8a). However, clonal expansions in recovered patients (R 1-3) were associated with specific mutations, except for a few clonotypes that duplicated without mutations. The coefficient of variation for each IGHV gene was found to be higher in recovered patients (Figure 8c).

Statistical analysis revealed significant differences in 26 out of 39 IGHV genes (one-tailed Mann-Whitney U-test, *p* < 0.05). This suggests a more diverse SHM pattern in recovered patients, potentially contributing to a more effective immune response for their recovery. In contrast, the deceased patients showed a limitation in BCR diversity due to less SHM, indicating a potential lack of effective immune response.

### Recovered patient BCR repertoires overlap with published COVID-19-specific BCRs

The antigen specificity and binding capacity of BCRs are determined by the sequences and structures of the CDR1, CDR2, and CDR3 regions on the heavy chain as well as light chain^35^, which we refer to here as the BCR paratopes. We compared our BCR data to published CDR datasets acquired from healthy or COVID-19 patient donors. In this analysis, we constructed a healthy database consisting of 223,405,489 BCRs from 15 studies of healthy donors comprising 330 repertoires (Table S15-1). We also constructed a COVID-19 database consisting of 8,882,096 BCRs from 18 studies of COVID-19 patients comprising 224 repertoires (Table S15-2). For each BCR in each patient in our cohort across two stages, we searched both the healthy database and the COVID-19 database to determine whether the number of hits was significantly more than expected (hypergeometric *p-*value < 0.001). We observed a total of 614 COVID-19 specific BCRs in our cohort, but only 4 apparently healthy specific BCRs, as expected (Table S16). Next, we assessed the COVID-19 specific BCRs as a fraction of each donor’s observed repertoire. The proportions of COVID-19 specific matches were higher in recovered patients than fatal patients (*p* = 0.0043, Figure 8d). These findings indicate that the B cell maturation process in recovered patients may be more effectively aligned towards producing BCRs capable of explicitly recognizing SARS-CoV-2.

## Discussion

It is likely that the viral strains affected the outcome of our patients, as the Alpha strain that infected all deceased patients was reported to have increased mortality of B.1.1.7-infected patients^36^. In Tokyo, Japan, B.1.1.7 strain was found to have higher transmissibility and S-protein mutations, but did not show a significant association of the Alpha strain with higher disease severity^37^. Several factors may have contributed to the outcome of the patients, including morphological changes in lymphocytes. In this study, such changes were observed in all patients (Table S17). Our COVID-19 patients show similar trends in prognostic clinical data to previous studies wherein critical cases showed an increase in white blood cell (WBC) counts, neutrophil counts, and neutrophil-lymphocyte ratio (NLR)^38^ as they progressed from the moderate stage. This suggests that all six patients reflected similar clinical status from the early moderate to the critical state. While it would have been ideal for all patients to have scRNA sequence data in all timepoints for more accurate comparison, the clinical data accounts for this limitation.

Using single-cell sequencing analysis of the PBMCs of Japanese patients, this study identified multiple factors to determine the prognosis of COVID-19. Clonal diversity after SARS-CoV-2 infection was indiscriminate in T cells but prognostically favorable in B cell clusters. Inefficient clonal expansion of B cells is prognostically poor, and differences in gene expression during critical illness—activation of biosynthetic processes in surviving patients and shift towards increased metabolism pathways in deceased patients—were highlighted.

The proliferation of CD4 TCM cells in both healthy subjects and COVID-19 patients highlights their protective role in developing viral infection^39, 40^. The abundant V-J gene regions (TRBV20-1, TRBV5-1, and TRBV19) were previously identified in recovered patients and in pre-pandemic samples^4, 41^, suggesting that pre-existing memory CD4 cells may have been reactivated upon COVID-19 infection.

The top 10 clones or the top 50% of the most abundant clones have been used to represent TCR repertoire diversity^42^. However, our data suggest that T cells in patients expanded twice as much as in healthy subjects (mean occupancy of COVID-19 patients = 0.4% of the total TCR repertoire), previous studies showed that large clones with occupancy above 0.5% are more correlated with positive response to treatment^43, 44^. The TCR repertoires obtained in our study lacked large clonal expansions but exhibited high diversity regardless of disease severity (seen in the % occupancy of all clonotypes). This diversity was not sufficient to protect the deceased patients, which is consistent with previous literature suggesting that TCR repertoire diversity is not correlated with effective viral clearance^45^.

Concerning BCRs, our study suggests that an expansion of clones above a threshold (>10 in the present study) is a marker of poor prognosis in COVID-19 infections (Figure S4). Despite a monoclonal expansion during the critical stage of the deceased patients, these antibodies failed to bind to known SARS-CoV-2 antigens in ELISA assays (Figure 8b and Figure S5). Monoclonal expansion exceeding four clone counts in the IGHV1-18 gene was observed in deceased patients during the critical period (D 1-3, Figure 8a and Figure S4). The use of the IGHV1-18 gene has been reported in acute dengue^46^ and proliferative lupus nephritis^47^, but not in COVID-19, suggesting the non-specificity of this receptor gene to SARS-CoV-2. IGHV4-34 was highly expanded in one of the deceased patients (D 2). This receptor was observed in COVID-19 patients in recovery in previous reports^45, 48^. However, in our cohort, it did not appear to be effective against the virus. Furthermore, increased usage of the IGHV genes was observed in patients who did not recover, suggesting a possible deviation in the immune response mechanism in these individuals.

This observation was also mirrored in the BCR reference repertoires, which indicated that the recovered patients demonstrated more hits to both healthy and COVID-19 BCRs than the patients who had fatal outcomes (Figure 8b and Table S16). We observed that a large proportion of private, or unshared clones were expressed in the BCR repertoire of the deceased patients. Sharing of receptors among individuals has been interpreted as evidence for common immune response to a pathogen^49, 50^, however, our results are insufficient to determine whether the shared BCR receptors in recovered patients were COVID-19-specific.

Notable variations in BCR subtypes were observed particularly in plasmablasts and naive cells. In recovered patients, the higher proportion of B naive cells may suggest that these patients had not yet undergone extensive B cell maturation or specialization. This could indicate that they retained a more diverse BCR repertoire, facilitating their recovery. Conversely, in patients who did not recover, a higher proportion of plasmablasts could imply a strong response to specific antigens or excessive activation of certain B cells. In contrast, there were no naïve CD4 cells recovered from all patients regardless of severity and outcome. Severe SARS-CoV-2 is notable for severe naïve T-cell depletion. Clones from the naive T-cell pool with high affinity to SARS-CoV-2 will differentiate into effector/memory T cells, resulting in a decrease in naive T cells^51^. Although a large pool of naïve T cells can determine the diversity of an activated T cell repertoire, it was irrelevant to our patients wherein naïve T cells are depleted even during the early stages of infection. The high frequency of naïve B cells during the critical stage, however, showed positive prognostics in the surviving patients.

These findings suggest that the repertoire diversity in T cells was independent of the presence of naïve cell, as well as disease severity and specific monoclonal immune responses in B cells to COVID-19 were ineffective against the infection, as opposed to the polyclonal B cell immune responses observed in survivors. Although cross-sectional comparisons were only performed on the critical state of all patients, it can be perceived that the repertoire of all patients during their early moderate state could be of similar conditions based on the similar trends of the peripheral blood cell quantity as summarized in the clinical data.

Furthermore, ineffective V(D)J recombination, clonal expansion and SHM observed in our cohort may indicate inadequacies in the underlying biological processes associated with the immune response to COVID-19. Activation of biological pathways based on gene expression detected in T and B cells during the critical phase differed significantly between deceased and surviving COVID-19 patients. During the critical period in surviving patients, macromolecular metabolic pathways (i.e., primary metabolic processes, nitrogen compound biosynthesis and protein metabolic processes) were activated in B and CD4 TCM cells. These pathways are involved in ribosome synthesis, translation and amino acid production. This finding is consistent with previous reports that activated immune cells increase ribosomal mRNA and produce large amounts of protein for proliferation and antiviral protein synthesis^52, 53^. On the other hand, expression of mitochondrial genes involved in ATP metabolism was increased in B and CD4 TCM cells during the critical period of the deceased patients. During cellular stress, mitochondria rapidly increase energy production and, in the absence of homeostatic mechanisms, induce an oversupply of oxidants such as reactive oxygen species^54^. Mitochondrial overactivity causes oxidative damage to cells, ultimately leading to loss of mitochondrial function. In a previous study, higher levels of oxidants and lower levels of antioxidants were detected in PBMCs of COVID-19 patients than in healthy controls^55^. Exhausted cells undergoing apoptosis also require a large quantity of ATP to ensure a smooth progression to apoptosis^56^. These findings suggest that mitochondrial dysfunction in immune cells during the critical phase is associated with poor prognosis.

Cell proliferation status with inflammatory signatures is known to hasten cell damage^57, 58^. We observed this during the critical state of the deceased cases, suggesting that persistent activation of inflammatory CD4 T cells may be associated with decreased survival in COVID-19 patients.

While the specific molecular mechanisms in B cells linking the V(D)J recombination, clonal expansions, and SHM processes to the broader biological pathway changes in COVID-19 remain unclear, it is crucial to consider these associations. The involvement of key enzymes such as recombination activating genes 1 and 2 (RAG)^59^, terminal deoxynucleotidyl transferase (TdT)^60, 61, 62^, Artemis nuclease^63^, Activation-induced cytidine deaminase (AID)^64^ in these processes underscores the complexity of the immune response. Further research is needed to clarify these connections and improve our understanding of the immune response to COVID-19.

This study has several limitations. First, this is a single center study with a small number of samples in Japan. Second, most of our patients were of advanced age, and immune responses may be different in younger patients. Third, other SARS-CoV-2 strains have emerged since the study period. Fourth, epitope prediction analysis was performed solely depending on the currently available public data, which could skew the results. Lastly, cross-sectional comparisons were only performed among critical samples between deceased and surviving patients. However, the similarity in the clinical data of all patients during their early moderate stages prior to progressing to critical can be a proxy for the lack of moderate samples of R2 and R3. Some discretion is warranted when considering applying our conclusions to patients with different strains.

In conclusion, this study highlights the importance of immune diversity and energy metabolism in COVID-19 outcome using single-cell repertoire analysis of cases in Japan, where the incidence and mortality rates were far lower than in other countries. Patients with different outcomes displayed different transcriptomic profiles in B and CD4 cells, with survivors exhibiting active biosynthetic pathways and deceased patients showing high expression of mitochondrial genes involved in energy production. Although clonal expansion patterns and repertoire diversity in T cells do not dictate the outcome of the infection, antigen-specific B cell receptors with a diverse repertoire can bolster immunity, as manifested in the survivors observed in this study. Also, surprisingly, a monoclonal expansion in B cells was associated with poorer outcome in COVID-19 patients. The diversity of the B cell response may be a key indicator of patient recovery, and this information may be of value when a new virus inevitably infects the human population.

## Materials and Methods

### Sample collection of PBMC for scRNA-seq

A total of 15 peripheral blood mononuclear cell (PBMC) samples were analysed. Twelve samples were collected from six COVID-19 inpatients admitted to Juntendo University Hospital in Tokyo, Japan. SARS-CoV-2 infection was established through RT-PCR-based molecular testing of collected nasopharyngeal specimens^65^ using the 2019 Novel Coronavirus Detection Kit (Shimadzu, Kyoto, Japan)^66^ at several points in time during different stages of disease progression (Table 1). Three samples were collected from three healthy subjects tested negative for COVID-19.

All patients received appropriate treatment during hospitalization, except for the deceased patient D 1, who refused ventilation and died 16 days from onset. Blood samples of D 1-3 and R 1 were collected during the moderate and the critical stage, while samples of R 2 and 3 were collected as they recovered from critical to moderate stage. D 1, 2, and 3 are deceased cases, and R 1, 2, and 3 are surviving cases. We categorized SARS-CoV-2 infection patients as moderate or critical according to the WHO criteria^67^. Moderate COVID-19 was defined as respiratory symptoms with clinical and radiological evidence of pneumonia. In cases, SpO2 ≥ 4% was observed on room air, while one of the following was required to identify the severe and critical cases: respiratory rate >30 breaths/min or SpO2 <94% on room air. Critical illness was defined as respiratory failure, septic shock, and/or multiple organ dysfunction (COVID-19 Clinical management: living guidance^67^). This study complied with all relevant national regulations and institutional policies. It was conducted in accordance with the tenets of the Declaration of Helsinki and was approved by the Institutional Review Board (IRB) at Juntendo University Hospital (IRB #20-037, #20-051). Written, informed consent was obtained from all individuals included in this study.

### PBMC preparation for single-cell suspension

PBMCs were isolated from blood samples using BD Vacutainer CPT mononuclear cell preparation tubes containing sodium heparin (BD Bioscience), following the manufacturer’s instructions. The isolated cells were washed with Dalbeccos’ phosphate buffered saline (PBS), resuspended in 1 mL of CELLBANKER 1 Plus, and stored at −80°C until further use.

### ScRNA-seq (10x platform)

PBMCs were rapidly thawed at 37°C and added RPMI-1640 containing heat-inactivated 10% v/v fetal bovine serum (FBS). The cells were confirmed to have over 90% viability via Trypan Blue staining. Cells with a concentration of around 700 – 1200 cells/uL were suspended in PBS containing 0.5% bovine serum albumin (BSA) and prepared to simultaneously generate paired V(D)J sequences and gene expression profiles using the 10x Chromium Single Cell platform using Chromium Next GEM single-cell 5’ GEM reagent kits v2 (1000263, 10x Genomics) according to the manufacturer’s protocol. The loaded cells were approximately 1.6 × 10^4^, aiming for 1.0 × 10^4^ single cells per reaction on the Chip K Single Cell Kit (1000286) with the Chromium controller (10x Genomics). Each cell was encapsulated in a microdroplet, which also contained a gel bead adorned with several oligonucleotide tags. Each of these oligonucleotide tags bore a beads-specific 10x barcode sequence (referred to as the cell-associated barcode) and was linked to one of various Unique Molecular Identifiers (UMI) sequences. Full-length cDNAs, tagged with the aforementioned oligonucleotide, were synthesized in the microdroplet. The cDNA were amplified and utilized for sequencing the V(D)J segments using the TCR Amplification Kit (1000252) and the BCR Amplification Kit (1000253), respectively.

ScRNA libraries were constructed with library-associated unique indexes, leveraging the library construction kit (1000196), the Single Index Kit T Set A (1000213), and SI primer (2000095). If necessary, these libraries underwent conversion using a Universal Library Conversion Kit (APP-A, 1000004155, MGI, China). The amplified cDNAs were cleaned and size selected using the SPRIselect magnetic beads (SPRIselect, Beckman-Coulter).

Quantification of both the amplifications and libraries was performed using the Agilent Bioanalyzer High Sensitivity DNA assay (High-Sensitivity DNA Kit, Agilent) and KAPA Library Quantification Kit (Kapa biosynthesis). This quantification was further confirmed through qRT-PCR, following the manufacturer’s instructions. The V(D)J and 5’ gene expression (GEX) libraries were mixed with the intention of achieving specific sequencing depths: 10,000 for TCR, 10,000 for BCR, and 30,000 for GEX libraries. The pooling ratios for each participant, as established based on a prior study by Stephenson and colleagues^18^, were as follows: 30:5:1 for COVID-19 critical cases, 12:2:1 for moderate cases, and 40:8:1 for healthy control. Sequencing was carried out on either the NovaSeq 6000 platform (Illumina) using a read length of 150-base paired-end or the DNBSEQ-G400 platform (MGI) with a 100-base paired-end read length. For gene expression, a minimum of 30,000 paired-end reads per cell was consistently achieved. Similarly, both TCR- and BCR-enriched libraries collected at least 10,000 paired-end reads per cell.

### Sequence alignment and assembly

Raw sequencing reads in FASTQ format were analyzed using the Cell Ranger (10x Genomics, version 6.1.2) multi pipeline, performing gene expression (GEX) alignment against the GRCh38 human reference dataset with a SARS-CoV-2 genomic RNA (GenBank: LC606022.1) appended. V(D)J assembly and alignment and paired clonotype calling for T and B cell receptors were done against the refdata-cellranger-vdj-GRCh38-alts-ensembl-5.0.0 reference (10x Genomics). CDR sequences and rearranged full-length BCR/TCR V(D)J segments as well as clonotype frequency were also obtained.

### scRNA-seq data pre-processing

Downstream scRNA-seq data analysis of the resulting expression count matrices from Cell Ranger was performed using the R programming language (version 4.2.1)^68^ with the Seurat package (version 4.0)^69^. All samples were integrated into one dataset, categorized according to severity and patient identity. Cell cycle phases were determined for each cell, which were judged to have no discernible effect on the variation of the data. Quality assessment and filtration methods include cell cycle phases, cells > 500 transcripts, > 250 genes, and < 20% mitochondrial genes. Genes with zero counts were also removed from the dataset. After this filtration, 129,469 cells were kept for further analysis. The data were normalized, and variable features were determined for clustering the cells. The Azimuth web application was used for cluster annotation^69^, and UMAP dimensionality reduction was used to view the resulting cell clusters/types. To account for interpatient variability, subsequent analyses were performed comparing individual patients.

### T and B Cell Repertoire Analysis

Gene expression data matrices obtained from the scRNA-seq analysis were processed to separately extract data for T cells and B cells. Filtered clonotype contigs from the TCR- and BCR-sequence assemblies were merged with the scRNA-seq data for each library using the *CombineExpression* function in the scRepertoire R package (version 1.7.0)^70^. Only cells with α- or β-, or both TCR chains, and light and heavy chains for BCR, were included in the repertoire dataset. In subsequent analyses for the TCR, we only investigated the β-chain region. TCR/BCR repertoire analysis was performed using the Seurat and scRepertoire packages in R.

### TCR clonotype antigen specificity and clonotype occupancy

We recovered transcripts encoding for the TCR β-chain from the TCR library to describe the clonal dynamics of the repertoire upon infection. At the onset of the disease, the CDR, specifically the CDR3, encoded in both the α- and β-chains, directly interact with the peptide antigen for viral clearance. For this reason, CDR3 is often used as the region of interest to determine T cell clonotypes, as it is highly unlikely that two T cells will express the same CDR3 nucleotide sequence, unless they have derived from the same clonally expanded T cell^71^. The β-chain, however, is known to be the more stable among the receptor chains of a TCR^72^. We then defined clonotypes based on the CDR3 sequences of the TCR β-chain using the Cell Ranger analysis pipeline.

The amino acid sequences of the CDR3β region of the TCR were used to predict the specificity to a known antigen referenced in four public databases (McPAS^29^, vdjdb^30^, TCRex^31^, and TCRMatch^32^), following criteria that CDR3 sequences should be long enough and should not differ by more than one amino acid as described by Gantner et al.^73^ and Meysman et al.^74^. A conservative threshold of 97-100% sequence similarity to reference sequences was observed and considered of high probability of sharing specificity.

### BCR CDRs Sequences overlap in Immunoglobulin Repertoire

Reference CDRs amino acid sequences of BCR from COVID-19 and healthy donors were prepared and used as databases for paratope-level similarity searching, as described previously^75^. In brief, we characterized each BCR’s heavy- and light-chain amino acid sequence by the three CDRs. A reference BCR was considered a match to a query if CDRs 1 and 2 both had sequence identities of 90% or more and CDR3 had a sequence identity of 80% or more. The pairwise repertoire overlap was given by the number of such matching BCRs, with significance given by hypergeometric p-value. These calculations were performed using an in-house python script (https://gitlab.com/sysimm/repoverlap).

### Analysis of Somatic Hypermutation and Clonal Expansion in BCR

BCR transcripts were annotated utilizing the IMGT/HighV-QUEST tool referencing the human immunoglobulin data set, available at https://imgt.org/IMGT_vquest^34^. Our primary assessment emphasized the V-gene usage along with the frequency of nucleotide mutations found within the variable region of the heavy chain. From this rigorous analysis, we derived data lists highlighting the “V-GENE and allele”, “V-REGION Number of nucleotides”, and “V-REGION Number of mutations”. The coefficient of variations for SHM ratio was calculated as follows: first, the SHM ratio was calculated for each clonotype. Then for each HV gene usage, the average SHM ratio and its standard deviation were determined. The coefficient of variation was obtained by dividing the standard deviation by the average SHM ratio.

### Differential gene expression and gene-set enrichment analysis

To identify the genes differentially expressed during each state for each patient, the data obtained at moderate stage were compared to those obtained at critical stage using the Wilcoxon test (fold-change of >1.5; adjusted p-value < 0.05), by the Seurat function *FindAllMarkers* (Bonferroni correction) for T cells, or the *wilcoxauc* (permutation testing) function from the presto package for B cells^76^. Gene set enrichment analysis was performed using the Hallmark functional gene set^77^.

Differential gene expression analysis was also performed between similar severity stages in different patient classifications—deceased vs. surviving in the critical stage and deteriorating vs. improving in the moderate stage. The identified marker genes were input into g:Profiler for functional profiling queries (g:GOSt) against the GO biological process gene ontology^78^. The results were mapped to enrichment networks in Cytoscape^26^ to visualize pathway overlap between the different patients in corresponding severity stages, and pathway clusters were generated using the Markov Cluster (MCL) algorithm^79^ to infer the relationships between the pathways. To quantify the UMI values and compare them among samples, the total UMI counts for each cell-associated barcode in the critical state of each patient were normalized towards the grand median UMI counts per sample library by a scaling factor (computed as total UMI counts per barcode/median UMI per sample library), as discussed in the 10X Genomics knowledge base. The cells were mapped based on the total UMI counts to identify the cell subtype as annotated in Azimuth. The cell clusters were then subset for differential gene expression analysis. Only libraries from the critical state of each patient were used.

### ELISA binding assay

#### 1. Cell lines and viruses

293T cells (RCB2202, RIKEN Cell Bank) and VeroE6/TMPRSS2 cells (JCRB1819, Japanese Collection of Research Bioresources Cell Bank) were maintained in DMEM supplemented with 10% FBS, penicillin (100 U/mL, Gibco), and streptomycin (100 μg/mL, Nacalai) and cultured at 37°C in 5% CO_2_. Expi293F cells (A14527, Thermo) were maintained in Expi293 expression medium (Gibco) supplemented with penicillin and streptomycin at 37℃ incubators under 8% CO_2_ and shaking at 125 rpm. The cells were routinely checked for mycoplasma contamination. SARS-CoV-2 was obtained from the National Institute for Infectious Diseases (WK-521 strain, GISAID ID: EPI_ISL_408667). The stock virus was amplified in VeroE6/TMPRSS2 cells. SARS-CoV-2 infection was carried out in a Biosafety Level 3 laboratory.

#### 2. Production of recombinant antibodies from COVID-19 patients-derived B cell clones

We produced recombinant anti-spike RBD antibodies and analyzed their binding capacity with specific proteins. Recombinant antibodies were produced as previously described^75^. Briefly, the variable regions of sampled heavy and light chains from the COVID-19 patients were prepared by dsDNA synthesis (IDT) and cloned into pCAGGS vectors containing sequences of human IgG1 or human kappa constant region, respectively. To produce recombinant antibodies, vectors containing heavy chain sequence and light chain sequence from known antibodies were co-transfected into Expi293F cells the day after pre-transfection seeding. The supernatants at day three were collected for further assay.

#### 3. Antibody binding to SARS-CoV-2 infected lysate

293T cells stably transfected with ACE2 and TMPRSS2 were prepared as previously reported (referred to as 293T-hACE2/hTMPRSS2)76. Either VeroE6/TMPRSS2 cells or 293T-hACE2/hTMPRSS2 cells were seeded at a density of 0.3×106 cells/mL in a 96-well plate and cultured for 18 hours. These cells were then infected with SARS-CoV-2 at a Multiplicity of Infection (MOI) of 0.05. At 36 hours post-infection, the cells were washed with PBS and resuspended in PBS containing 0.1% Triton X, 2% CHAPS, and a protease inhibitor. Mixtures from 293T-hACE2/hTMPRSS2 and VeroE6/TMPRSS2 cell mock or SARS-CoV-2-infected were centrifuged at 3000 rpm for 5 minutes and 4℃. The supernatant was discarded, and the pellet was vortexed resulting in around 200 µL of resuspended cell pellet. The cell pellet was coated at 50 times dilution with PBS onto half-area 96-wells plate and incubated overnight at 4℃. After two washes with PBS, the plate was blocked with 1% BSA in PBS. Following two washes, plates were then incubated with 25 *μ*l/well of a 1 *μ*g/ml solution of recombinant antibody or control antibodies (anti-PTx, Pertussis Toxinm, Ab49, in-house; Anti SARS-CoV-2 S RBD, 40592-R001, Sino Biological) was applied and incubated for 2 hours at room temperature. After three times washing, HRP-conjugated anti-human IgG (109-036-003, Jackson ImmunoResearch) or HRP-conjugated anti-Rabbit IgG (4030-05, SouthernBiotech) were added and incubated for 40 minutes. After two washes, substrate in TMBZ (Bakelite Sumitomo) was added to each well for 10 minutes and the reaction was stopped using H2SO4 solution (Sumitomo Bakelite). The experiments were performed in duplicates and the OD450 was measured using GloMax Explorer Multimode Microplate Reader (Promega).

### Statistical analysis

The Shannon index was used to calculate V gene usage and Ig isotype diversity scores for the individual samples^80^, and Wilcoxon’s test was used to compare the clonal and isotype diversity indices between samples, to compare isotype proportions, and to detect differentially expressed genes (DEGs) between different subsets of the data^81^. In all statistical comparisons, p values less than 0.05 were considered significant. All statistical analyses were performed using the R programming language^68^.

## Acknowledgments

We would like to sincerely thank all the persons involved in investigating of the SARS-CoV-2 infections and the patients at Juntendo University Hospital, Japan. We thank Ms. Kaori Saito for her help with scRNA seq. We are grateful to Dr. Shuhei Sakakibara and Dr. Yoshiaki Nishiya for their invaluable support in informatics. This study was conducted with the support of the Section of Diagnostics and Therapeutics of Intractable Diseases of Intractable Disease Research Center and Departments of Research Support Utilizing Bioresource Bank and Metabolism and Endocrinology, Juntendo University Graduate School of Medicine. This work was partially supported by the Japan Agency for Medical Research and Development (JP20fk0108472 to TN), by Japan Society for the Promotion of Science Grants-in Aid for Scientific Research (22K08608 to MS and 22K15675 to ST) and by Research Support Project for Life Science and Drug Discovery (Basis for Supporting Innovative Drug Discovery and Life Science Research (BINDS)) from AMED (JP22ama121025 to DS).

## Contributions

Y.T., T.A., T.N., D.M.S. conceived and designed the study; R.O., F.J.P., K.Y. performed the single cell RNA seq experiments; C.O. conducted antibody binding experiments; K.T. directed antibody binding experiments; A.K., F.J.P., A.A.S., D.S.S., H.S.I., R.O. conducted data analysis; F.J.P., A.K., T.A., D.M.S., D.S.S., Y.T., R.O., A.A.S. wrote the paper; M.S., T.N., S.T., Y.H., K.T. took care of patients and provided the clinical information; F.J.P., A.K., T.A., R.O., Y.T., D.M.S. provided intellectual input throughout the study, provided comments and helped edit the paper. All authors read and approved the final paper.

## Competing interests

All authors declare that there are no competing interests.

## Data availability

Raw and processed data used in this study are available on NCBI’s Gene Expression Omnibus (GEO) database.^82^ The files are accessible through the following accession numbers: GEX, GSE267645 (https://www.ncbi.nlm.nih.gov/geo/query/acc.cgi?acc=GSE267645); TCR, GSE267639 (https://www.ncbi.nlm.nih.gov/geo/query/acc.cgi?acc=GSE267639); BCR, GSE267642 (https://www.ncbi.nlm.nih.gov/geo/query/acc.cgi?acc=GSE267642).

## Supplementary information

**Figure S1. Percentage occupancy of top 50 most abundant clonotypes.** Clonotypes with predicted specificity to SARS-COV-2 are shown in red dots. The proportions of the mean occupancy of the Top 50 clonotypes in the whole TCR repertoire and the occupancy of all clonotypes are presented.

**Figure S2. V and J gene usage for T cells.** V genes are shown in the x-axis; bar heights represent V gene percentage in each sample; colors represent J genes associated with respective V genes in CD4 TCM **(a)**, and CD8 TEM **(b)**.

**Figure S3. V and J gene usage for B cells.** V genes are shown in the x-axis; bar heights represent V gene percentage in each sample; colors represent J genes associated with respective V genes. The scale for the y-axis in COVID-19 samples is 0-20%, except for Pt 2, where, in the critical sample, the IGHV4-34 gene occupies over 40% of the V gene repertoire. Survivors show similar V–J combination proportions to healthy individuals, whereas deceased patients show clonal expansion patterns that are patient-specific, most pronounced in Pt 2’s critical sample.

**Figure S4.** Clonotypes illustrated with somatic hypermutation (SHM) ratio and clonal expansion in relation to IGHV gene usage in critical stage. Each clonotype is represented as bubble with the size indicating the count of each clonotype. The position of each bubble corresponds to the specific HV gene usage and the SHM rate. Arrows in the figure highlight those expanded clonotypes that were selected for subsequent antibody binding assays, indicating their importance in the context of the study.

**Figure S5.** Antibody binding assay against lysate from SARS-CoV-2 infected cells. Recombinant antibodies were generated using sequences from recovered patients (R 2 and R3) and deceased patients (D 1 and D 2). VeroE6/TMPRSS2 and 293T cells were seeded in equal numbers and cultured for 15 hrs. before being infected with SARS-CoV-2 for an additional 6 hrs. Lysates from these infected cells were then immobilized on ELISA plates, followed by incubation with the recombinant antibodies. Antibody binding was detected using HRP-labeled anti-human IgG. Cells not infected with the virus showed greater proliferation, leading to enhanced antibody binding as seen with anti-beta actin antibodies. Notably, higher binding compared to the non-infected samples was observed in p5_3 and in anti-S RBD antibody. These assays were performed in duplicate.

**Figure S6.** Analysis of Somatic Hypermutations (SHM) in BCR Clonotypes during the Critical Stage of COVID-19. The heavy chain sequences of BCR clone were analyzed the SHM using IMGT/HighV-Quest. We visualized the aggregate number of mutations and the percentage of clonotypes with mutations for each sample using tables and bar graphs. The deceased patients demonstrated more mutations and a greater percentage of mutated clonotypes than others (Welch’s t-test, p = 0.041 or 0.016).

## Table Legends

Table S1. Frequency of T cell subtypes.

Table S2. Differential gene expression for each CD4 TCM TRBV subset in moderate vs critical states.

Table S3. Hallmark biological pathways in CD4 TCM cells during the moderate state.

Table S4. Upregulated gene expressions during the critical states of deceased (D 1-3) vs. surviving (R 1-3) patients in CD4 TCM TRBV subset.

Table S5. Gene ontology biological pathways of upregulated genes in CD4 TCM TRBV20-1 subset during critical states of deceased (D 1-3) vs surviving patients (R 1-3).

Table S6. Upregulated gene expressions during the critical states of deceased (D 1-3) vs. surviving (R 1-3) patients in CD4 proliferating cell subset.

Table S7. Antigen affinity prediction of T cell CDR3β clonotypes.

Table S8. Frequency of V-gene usage in CD4 TCM cells.

Table S9. Frequency of V-gene usage in CD8 TEM cells.

Table S10. Frequency of B cell subtypes.

Table S11. Differential gene expression of B cells in moderate vs critical states.

Table S12. Hallmark biological pathways in B cells during the moderate state.

Table S13. Gene ontology biological pathways of upregulated genes in B cells during critical states of deceased (D 1-3) vs surviving patients (R 1-3).

Table S14. Frequency of V-gene usage in B cells.

Table S15. BCR repertoire sequence public databases.

Table S16. BCR overlaps with previous studies.

Table S17. Blood cell counts of patients with COVID-19 during different time points.

## Notes

### Competing Interest Statement

The authors have declared no competing interest.

### Summary of Updates

Figure layout updated to show previously hidden parts

## References

1. Chen Z, John Wherry E. T cell responses in patients with COVID-19. Nature Reviews Immunology 20, 529–536 (2020).

2. Huang C, et al. Clinical features of patients infected with 2019 novel coronavirus in Wuhan, China. Lancet 395, 497–506 (2020).

3. Lucas C, et al. Longitudinal analyses reveal immunological misfiring in severe COVID-19. Nature 584, 463–469 (2020).

4. Zhang F, et al. Adaptive immune responses to SARS-CoV-2 infection in severe versus mild individuals. Signal Transduct Target Ther 5, 156 (2020).

5. COMBAT. A blood atlas of COVID-19 defines hallmarks of disease severity and specificity. Cell 185, 916–938.e958 (2022).

6. Masood KI, et al. Upregulated type I interferon responses in asymptomatic COVID-19 infection are associated with improved clinical outcome. Sci Rep-Uk 11, 22958 (2021).

7. Hadjadj J, et al. Impaired type I interferon activity and inflammatory responses in severe COVID-19 patients. Science 369, 718–724 (2020).

8. Luo L, et al. Dynamics of TCR repertoire and T cell function in COVID-19 convalescent individuals. Cell Discov 7, 89 (2021).

9. Ambikan AT, et al. Multi-omics personalized network analyses highlight progressive disruption of central metabolism associated with COVID-19 severity. Cell Syst 13, 665–681 e664 (2022).

10. Nunn AVW, et al. SARS-CoV-2 and mitochondrial health: implications of lifestyle and ageing. Immunity & Ageing 17, 33 (2020).

11. Ramesh S, Park S, Im W, Call MJ, Call ME. T cell and B cell antigen receptors share a conserved core transmembrane structure. Proc Natl Acad Sci U S A 119, e2208058119 (2022).

12. Jacob J, Kelsoe G, Rajewsky K, Weiss U. Intraclonal generation of antibody mutants in germinal centres. Nature 354, 389–392 (1991).

13. Saputri DS, et al. Deciphering the antigen specificities of antibodies by clustering their complementarity determining region sequences. mSystems, e0072223 (2023).

14. Chang CM, Feng PH, Wu TH, Alachkar H, Lee KY, Chang WC. Profiling of T Cell Repertoire in SARS-CoV-2-Infected COVID-19 Patients Between Mild Disease and Pneumonia. J Clin Immunol 41, 1131–1145 (2021).

15. Cao Y, et al. Potent Neutralizing Antibodies against SARS-CoV-2 Identified by High-Throughput Single-Cell Sequencing of Convalescent Patients’ B Cells. Cell 182, 73–84 e16 (2020).

16. Bacher P, et al. Low-Avidity CD4(+) T Cell Responses to SARS-CoV-2 in Unexposed Individuals and Humans with Severe COVID-19. Immunity 53, 1258–1271.e1255 (2020).

17. Meckiff BJ, et al. Imbalance of Regulatory and Cytotoxic SARS-CoV-2-Reactive CD4(+) T Cells in COVID-19. Cell 183, 1340–1353.e1316 (2020).

18. Stephenson E, et al. Single-cell multi-omics analysis of the immune response in COVID-19. Nat Med 27, 904–916 (2021).

19. Liao M, et al. Single-cell landscape of bronchoalveolar immune cells in patients with COVID-19. Nat Med 26, 842–844 (2020).

20. Liu C, et al. Time-resolved systems immunology reveals a late juncture linked to fatal COVID-19. Cell 184, 1836–1857.e1822 (2021).

21. Bernardes JP, et al. Longitudinal Multi-omics Analyses Identify Responses of Megakaryocytes, Erythroid Cells, and Plasmablasts as Hallmarks of Severe COVID-19. Immunity 53, 1296–1314.e1299 (2020).

22. Bieberich F, et al. A Single-Cell Atlas of Lymphocyte Adaptive Immune Repertoires and Transcriptomes Reveals Age-Related Differences in Convalescent COVID-19 Patients. Front Immunol 12, 701085 (2021).

23. He B, et al. Rapid isolation and immune profiling of SARS-CoV-2 specific memory B cell in convalescent COVID-19 patients via LIBRA-seq. Signal Transduct Target Ther 6, 195 (2021).

24. Jin X, et al. Global characterization of B cell receptor repertoire in COVID-19 patients by single-cell V(D)J sequencing. Brief Bioinform 22, (2021).

25. Khare S, et al. GISAID’s Role in Pandemic Response. China CDC Wkly 3, 1049–1051 (2021).

26. Shannon P, et al. Cytoscape: a software environment for integrated models of biomolecular interaction networks. Genome Res 13, 2498–2504 (2003).

27. Bonyek-Silva I, et al. LTB(4)-Driven Inflammation and Increased Expression of ALOX5/ACE2 During Severe COVID-19 in Individuals With Diabetes. Diabetes 70, 2120–2130 (2021).

28. Ren X, et al. COVID-19 immune features revealed by a large-scale single-cell transcriptome atlas. Cell 184, 1895–1913.e1819 (2021).

29. Tickotsky N, Sagiv T, Prilusky J, Shifrut E, Friedman N. McPAS-TCR: a manually curated catalogue of pathology-associated T cell receptor sequences. Bioinformatics 33, 2924–2929 (2017).

30. Shugay M, et al. VDJdb: a curated database of T-cell receptor sequences with known antigen specificity. Nucleic Acids Res 46, D419–d427 (2018).

31. Gielis S, et al. Detection of Enriched T Cell Epitope Specificity in Full T Cell Receptor Sequence Repertoires. Frontiers in Immunology 10, (2019).

32. Chronister WD, et al. TCRMatch: Predicting T-Cell Receptor Specificity Based on Sequence Similarity to Previously Characterized Receptors. Frontiers in Immunology 12, (2021).

33. Rajewsky K. Clonal selection and learning in the antibody system. Nature 381, 751–758 (1996).

34. Alamyar E, Giudicelli V, Duroux P, Lefranc M-P. Antibody V and C Domain Sequence, Structure, and Interaction Analysis with Special Reference to IMGT®. In: Monoclonal Antibodies: Methods and Protocols (eds Ossipow V, Fischer N). Humana Press (2014).

35. Chitarra V, et al. Three-dimensional structure of a heteroclitic antigen-antibody cross-reaction complex. Proc Natl Acad Sci U S A 90, 7711–7715 (1993).

36. Davies NG, et al. Increased mortality in community-tested cases of SARS-CoV-2 lineage B.1.1.7. Nature 593, 270–274 (2021).

37. Tsuchiya K, et al. Molecular characterization of SARS-CoV-2 detected in Tokyo, Japan during five waves: Identification of the amino acid substitutions associated with transmissibility and severity. Frontiers in Microbiology 13, (2022).

38. Horiuchi Y, et al. Peripheral granular lymphocytopenia and dysmorphic leukocytosis as simple prognostic markers in COVID-19. International Journal of Laboratory Hematology 43, 1309–1318 (2021).

39. Simnica D, et al. Landscape of T-cell repertoires with public COVID-19-associated T-cell receptors in pre-pandemic risk cohorts. Clin Transl Immunology 10, e1340 (2021).

40 . Mahajan S, et al. Immunodominant T-cell epitopes from the SARS-CoV-2 spike antigen reveal robust pre-existing T-cell immunity in unexposed individuals. Scientific Reports 11, 13164 (2021).

41. Wu D, et al. Structural assessment of HLA-A2-restricted SARS-CoV-2 spike epitopes recognized by public and private T-cell receptors. Nat Commun 13, 19 (2022).

42. Hosoi A, et al. Author Correction: Increased diversity with reduced “diversity evenness” of tumor infiltrating T-cells for the successful cancer immunotherapy. Sci Rep-Uk 13, 6816 (2023).

43. Fairfax BP, et al. Peripheral CD8(+) T cell characteristics associated with durable responses to immune checkpoint blockade in patients with metastatic melanoma. Nat Med 26, 193–199 (2020).

44. Kidman J, et al. Characteristics of TCR Repertoire Associated With Successful Immune Checkpoint Therapy Responses. Front Immunol 11, 587014 (2020).

45. La Gruta NL, et al. Epitope-specific TCRbeta repertoire diversity imparts no functional advantage on the CD8+ T cell response to cognate viral peptides. Proc Natl Acad Sci U S A 105, 2034–2039 (2008).

46. Godoy-Lozano EE, et al. Lower IgG somatic hypermutation rates during acute dengue virus infection is compatible with a germinal center-independent B cell response. Genome Med 8, 23 (2016).

47. Ghasemi M, et al. Predictive Biomarker Panel in Proliferative Lupus Nephritis-Two-Dimensional Shotgun Proteomics. Iran J Kidney Dis 1, 121–133 (2021).

48. Zhang F, et al. Correction to: Adaptive immune responses to SARS-CoV-2 infection in severe versus mild individuals. Signal Transduct Target Ther 6, 161 (2021).

49. Galson JD, et al. Deep Sequencing of B Cell Receptor Repertoires From COVID-19 Patients Reveals Strong Convergent Immune Signatures. Front Immunol 11, 605170 (2020).

50. Nielsen SCA, et al. Human B Cell Clonal Expansion and Convergent Antibody Responses to SARS-CoV-2. Cell Host Microbe 28, 516–525.e515 (2020).

51. Zhang W, Zhou Y, Kang Y. Naïve T cells may be key to the low mortality of children with COVID-19. Journal of Evidence-Based Medicine 15, 3–5 (2022).

52. Wolf T, et al. Dynamics in protein translation sustaining T cell preparedness. Nature Immunology 21, 927–937 (2020).

53. Araki K, et al. Translation is actively regulated during the differentiation of CD8+ effector T cells. Nature Immunology 18, 1046–1057 (2017).

54. Eisner V, Picard M, Hajnóczky G. Mitochondrial dynamics in adaptive and maladaptive cellular stress responses. Nature Cell Biology 20, 755–765 (2018).

55. Tavassolifar MJ, et al. New insights into extracellular and intracellular redox status in COVID-19 patients. Redox Biology 59, 102563 (2023).

56. Chen LY, Yang B, Zhou L, Ren F, Duan ZP, Ma YJ. Promotion of mitochondrial energy metabolism during hepatocyte apoptosis in a rat model of acute liver failure. Mol Med Rep 12, 5035–5041 (2015).

57. Kiraly O, Gong G, Olipitz W, Muthupalani S, Engelward BP. Inflammation-induced cell proliferation potentiates DNA damage-induced mutations in vivo. PLoS Genet 11, e1004901 (2015).

58. Sacconi A, et al. Multi-omic approach identifies a transcriptional network coupling innate immune response to proliferation in the blood of COVID-19 cancer patients. Cell Death Dis 12, 1019 (2021).

59. Jones JM, Gellert M. The taming of a transposon: V(D)J recombination and the immune system. Immunol Rev 200, 233–248 (2004).

60. Landau NR, St John TP, Weissman IL, Wolf SC, Silverstone AE, Baltimore D. Cloning of terminal transferase cDNA by antibody screening. Proc Natl Acad Sci U S A 81, 5836–5840 (1984).

61. Isobe M, et al. Chromosome localization of the gene for human terminal deoxynucleotidyltransferase to region 10q23-q25. Proc Natl Acad Sci U S A 82, 5836–5840 (1985).

62. Yang-Feng TL, Landau NR, Baltimore D, Francke U. The terminal deoxynucleotidyltransferase gene is located on human chromosome 10 (10q23----q24) and on mouse chromosome 19. Cytogenet Cell Genet 43, 121–126 (1986).

63. Moshous D, et al. Artemis, a novel DNA double-strand break repair/V(D)J recombination protein, is mutated in human severe combined immune deficiency. Cell 105, 177–186 (2001).

64. Muramatsu M, et al. Specific expression of activation-induced cytidine deaminase (AID), a novel member of the RNA-editing deaminase family in germinal center B cells. J Biol Chem 274, 18470–18476 (1999).

65. Jin B, et al. Investigation of the individual genetic evolution of SARS-CoV-2 in a small cluster during the rapid spread of the BF.5 lineage in Tokyo, Japan. Front Microbiol 14, 1229234 (2023).

66. Fukumoto T, et al. Efficacy of a novel SARS-CoV-2 detection kit without RNA extraction and purification. Int J Infect Dis 98, 16–17 (2020).

67. WHO. Living guidance for clinical management of COVID-19.) (2021).

68. R Core Team. R: A language and environment for statistical computing. R Foundation for Statistical Computing.). 4.2.1 edn (2022).

69. Hao Y, et al. Integrated analysis of multimodal single-cell data. Cell 184, 3573–3587.e3529 (2021).

70. Borcherding N. scRepertoire: A toolkit for single-cell immune receptor profiling.). 1.7.0 edn (2022).

71. Rosati E, Dowds CM, Liaskou E, Henriksen EKK, Karlsen TH, Franke A. Overview of methodologies for T-cell receptor repertoire analysis. BMC Biotechnol 17, 61 (2017).

72. Haga-Friedman A, Horovitz-Fried M, Cohen CJ. Incorporation of transmembrane hydrophobic mutations in the TCR enhance its surface expression and T cell functional avidity. J Immunol 188, 5538–5546 (2012).

73. Gantner P, et al. Single-cell TCR sequencing reveals phenotypically diverse clonally expanded cells harboring inducible HIV proviruses during ART. Nat Commun 11, 4089 (2020).

74. Meysman P, De Neuter N, Gielis S, Bui Thi D, Ogunjimi B, Laukens K. On the viability of unsupervised T-cell receptor sequence clustering for epitope preference. Bioinformatics 35, 1461–1468 (2019).

75. Ismanto HS, et al. Landscape of infection enhancing antibodies in COVID-19 and healthy donors. Computational and Structural Biotechnology Journal 20, 6033–6040 (2022).

76. Browning BL. PRESTO: Rapid calculation of order statistic distributions and multiple-testing adjusted P-values via permutation for one and two-stage genetic association studies. BMC Bioinformatics 9, 309 (2008).

77. Liberzon A, Birger C, Thorvaldsdóttir H, Ghandi M, Mesirov JP, Tamayo P. The Molecular Signatures Database (MSigDB) hallmark gene set collection. Cell Syst 1, 417–425 (2015).

78. Raudvere U, et al. g:Profiler: a web server for functional enrichment analysis and conversions of gene lists (2019 update). Nucleic Acids Research 47, W191–W198 (2019).

79. van Dongen S, Abreu-Goodger C. Using MCL to extract clusters from networks. Methods in molecular biology 804, 281–295 (2012).

80. Keylock CJ. Simpson diversity and the Shannon–Wiener index as special cases of a generalized entropy. Oikos 109, 203–207 (2005).

81. Wilcoxon F. Individual comparisons of grouped data by ranking methods. J Econ Entomol 39, 269 (1946).

82. Barrett T, et al. NCBI GEO: archive for functional genomics data sets—update. Nucleic Acids Research 41, D991–D995 (2012).

